# Reproductive strategies in a complex of simultaneously hermaphroditic species, the *Allolobophora chlorotica* case study

**DOI:** 10.1101/2022.01.31.475338

**Authors:** L. Dupont, H. Audusseau, D. Porco, K. R. Butt

**Affiliations:** Univ Paris-Est Creteil, CNRS, INRAE, IRD, IEES-Paris, F-94010 Creteil France; Sorbonne Université, IEES-Paris, F-75005 Paris, France; Université de Paris, IEES-Paris, F-75013 Paris, France; CNRS UMR 6553 ECOBIO, Université Rennes 1, Avenue Général Leclerc, 35000 Rennes, France; Musée national d’histoire naturelle, L-2160 Luxembourg; University of Central Lancashire, School of Natural Sciences, Preston, PR2 6AG, UK

**Keywords:** Earthworms, Hybridization, Mitochondrial lineage, Multiple mating, Parentage analysis, Post-zygotic isolation

## Abstract

An in-depth knowledge of reproductive strategies is essential to understand the evolutionary history of species and their resulting patterns of taxonomic diversity. In particular, the study of life history traits related to reproduction could help to resolve the speciation patterns in the cryptic species complexes recently found in earthworms. Here, we carried out a parentage analysis in such a complex, the *Allolobophora chlorotica* aggregate. Using four nuclear microsatellite markers and a fragment of the *cytochrome c oxidase subunit I* mitochondrial gene we investigated (i) the mating strategies between individuals belonging to two divergent mitochondrial lineages (L2 and L3) that cannot be distinguished with nuclear markers and (ii) the reproductive isolation between lineages that are differentiated both at the mitochondrial and nuclear level (L2/L3 and L1). Among the 157 field collected individuals, 66 adults were used in cross-breeding experiments to form 22 trios based on their assignment to a mitochondrial lineage, and 453 obtained juveniles were genotyped. We showed that adults that mated with both their potential mates in the trio produced significantly more juveniles. In L2 and L3 crosses, a sex-specific pattern of reproduction characteristic to each lineage was observed, suggesting a possible conflict of interest between mating partners. In L2/L3 and L1 crosses, a high production of cocoons was counterbalanced by a low hatching rate, suggesting a post-zygotic reproductive isolation. Reproductive strategies are thus likely to contributed to the diversification of this complex of species and we advocate further studies on sexual selection and sex allocation in earthworms.

## Introduction

The diversity of earthworm species has long been underestimated because works in systematics mostly relied solely on morphological characters. Over the past decades, studies that have applied molecular techniques to explore earthworm systematics, taxonomy, and phylogeography discovered an unprecedented cryptic diversity revealing the recurrent discordance between traditional (morphology-based) and molecular (DNA-based) methods for assessing earthworm diversity (e.g. King, Tibble and Symondson, 2008; Novo *et al*., 2010; Dupont *et al*., 2011; Taheri *et al*., 2018). The processes that generated this diversity remain poorly understood, although these studies have identified factors that may contribute to it (Marchán *et al*., 2018). These factors may be extrinsic, such as paleogeographical (Shekhovtsov *et al*., 2013; Marchán *et al*., 2017) and ecological (Novo *et al*., 2012; Spurgeon *et al*., 2016). Others are intrinsic and result, for example, from differences in life history traits and reproductive strategies among species (Jones *et al*., 2016; Marchán *et al*., 2017). In particular, reproductive strategies which are subject to many evolutionary forces, including natural and sexual selection, and sexual conflict (Anholt *et al*., 2020) are known to contribute to various patterns of reproductive isolation (e.g. Noh & Henry, 2010) and, consequently, on speciation processes and diversity patterns (Mayr, 1963). Indeed, in the absence of reproductive isolation, interbreeding between species should reduce the taxonomic diversity (Rabosky, 2016).

The *Allolobophora chlorotica* complex of earthworm species, which was previously described as an aggregate of several closely related lineages (Dupont *et al*., 2016), is a good model to study reproductive strategies and isolation. These lineages exist as two colour morphs, green and pink (Satchell, 1967; King *et al*., 2008; Dupont *et al*., 2016). The green morph represents a single species composed of two divergent mitochondrial lineages that cannot be distinguished with nuclear markers (i.e. L2 and L3). The taxonomic status of the pink morph is less clear, even though it is believed to be composed of at least three species corresponding to three mitochondrial lineages (i.e. L1, L4 and L5, Dupont *et al*., 2011; King *et al*., 2008). Field and laboratory observations showed that the green morph tends to be more common in wet soils and the pink morph in dry soils (Satchell, 1967; Lowe & Butt, 2007). Thus, although it is often stated that these lineages have a sympatric distribution (i.e they co-occur in a region), some of them at least do not live in syntopy (i.e. they do not use the same habitat, Rivas, 1964). These distinct ecological preferences led Lowe & Butt (2007) to suggest that soil moisture acts as a prezygotic barrier to intermorphic mating in syntopic populations of these simultaneous hermaphrodite earthworms that do not self-fertilized (Dupont *et al*., 2011). Breeding experiments additionally reported mechanisms of postzygotic isolation. The viability of cocoons resulting from the crossing between the two colour morphs is severely restricted and the male offspring from the backcrossing of hybrids with pure bred morphs are sterile (Lowe & Butt, 2008). The genotyping of individuals collected in two natural populations also supported the hypothesis of a process of reproductive isolation between L1 (pink morph) and L2/L3 (green morph) lineages (Dupont *et al*., 2016). Dupont *et al*. (2016) showed a high level of congruence between the assignment based on the mitochondrial COI and the nuclear microsatellites, suggesting that hybridization is uncommon between L1 and L2/L3. However, they also reported cases of introgression, with a few individuals having a L1 haplotype but assigned to a nuclear cluster grouping all the L2/L3 individuals (Dupont *et al*., 2016). These individuals could result from multiple generations of unidirectional hybridization between L1 females and L2/L3 males. Each backcross with L2/L3 would dilute the proportion of L1 nuclear allele by half until the population had overwhelmingly accumulated nuclear L2/L3 alleles but retained the maternally inherited L1 mtDNA haplotype. The authors also recorded one individual having a L2 haplotype but assigned to the nuclear cluster grouping the majority of the L1 individuals, suggesting that unidirectional hybridization could also happen from mating between L2/L3 females and L1 males (Dupont *et al*., 2016). These events of unidirectional hybridization reinforce the hypothesis of sterility of the male function in L1-L2/L3 crosses.

Here, we combined cross-breeding experiments and a parentage analysis based on the sequencing of the *cytochrome c oxidase subunit I* mitochondrial gene and microsatellite markers, to get insights into the reproductive strategy and the reproductive isolation within this intriguing complex of closely related earthworm lineages, the *A. chlorotica* aggregate. First, we investigated whether earthworms of this species complex may have offspring from multiple mates during the same mating period. Second, we examined mating strategies between L2 and L3, in order to detect any evidence of reproductive isolation between these divergent mitochondrial lineages which could not be differentiated by nuclear markers in previous studies (Dupont *et al*., 2011, 2016). Third, we tested the hypothesis that L1 and L2/L3 individuals preferentially mate with a partner of the same taxon. Finally, we examined whether one function (male or female) is preferentially used in mating between these lineages.

## Material and Methods

The work described below corresponds to the parentage analysis of the offspring of *A. chlorotica* individuals collected in the field. We carried out cross-breeding experiments based on the mitochondrial assignment of the collected individuals. To assign the parentage of their offspring, we used both nuclear (i.e. microsatellites) and mitochondrial markers. This genotyping, using microsatellite markers, made it possible to finely characterize the genetic structure of the field population.

### Field sampling and genetic analysis of collected specimens

The sampling was carried out at a farm site (Walton Hall Farm, Preston, UK) in a field grazed by cattle. The soil is an alluvial, sandy clay with a pH of 8.3. Regardless of their colour morph, we collected a total of 157 *A. chlorotica* individuals: 46 juveniles in 2012, and 71 juveniles and 40 adults (i.e. clitellate) in 2015. From each individual, we removed the last segments of the caudal section and preserved it in 96% ethanol before DNA extraction using the NucleoSpin^®^ 96 Tissue kit (Macherey-Nagel). It is worth noting that the ablation of the last segments of the caudal section of earthworms does not affect their survival due to their regeneration capacity (e.g. Xiao *et al*., 2011).

#### Assignment of the collected individuals to *A. chlorotica* mitochondrial lineages

To assign the collected individuals to *A. chlorotic*a mitochondrial lineages, we amplified and sequenced the fragment of the *cytochrome c oxidase subunit I* mitochondrial gene (COI), proposed as a standard DNA barcode for animals (Hebert *et al*., 2003). Individuals collected in the field population in 2012 were processed at the Canadian Centre for DNA barcoding (CCDB), in the context of the global earthworm barcoding campaign (EarthwormBOL, Rougerie *et al*., 2009) and as part of the International Barcode of Life Initiative (iBOL). PCR amplifications and DNA sequencing were performed according to the standard protocols used in CCDB (Hajibabaei *et al*., 2005). For the individuals collected in the field population in 2015, the COI gene was amplified using the primer pair described in Folmer *et al*. (1994). DNA sequencing was performed by Eurofins Genomics company and we manually aligned the sequences using the BioEdit program (Hall, 1999). We inferred the mitochondrial lineage of the individuals using the identification engine of BOLD (Barcode of Life Data Systems – https://www.boldsystems.org/).

#### Genetic structure of the field population

To explore the mitochondrial genetic variation in the field population, we estimated haplotype frequencies using the software DNASP 6.0 (Rozas *et al*., 2017). We then examined the relationships among haplotypes using a haplotypic network constructed using a reduced-median algorithm (Bandelt *et al*., 1995) as implemented in the software NETWORK 10 (www.fluxus-engineering.com/sharenet.htm).

To explore the nuclear genetic variation in the field population, we genotyped individuals at four highly polymorphic microsatellite loci, Ac127, Ac170, Ac418 and Ac476, as described in Dupont *et al*. (2011). We amplified the loci by polymerase chain reaction (PCR) in one multiplex set and in 12 μl reactions using 10 ng of DNA and the Qiagen ^®^ Multiplex Kit, according to the manufacturer’s protocol. The migration of the PCR products was carried out on an ABI 3130 xl Genetic Analyzer using the LIZ500 size standard (Applied Biosystems); alleles were scored using GeneMapper 5 software (Applied Biosystems). We then calculated all basic genetic parameters including allele frequencies, number of alleles (N_all_), polymorphic information Content (PIC) for each locus and the observed (H_o_) and expected (H_e_) heterozygosity using CERVUS v.3.0.7 (Marshall *et al*., 1998; Kalinowski *et al*., 2007). We tested for the null independence between loci from statistical genotypic disequilibrium analysis using Genepop V4.4 (Rousset, 2008). The significance of a deviation from the Hardy-Weinberg equilibrium including a Bonferroni correction and null allele frequencies were estimated using CERVUS v.3.0.7.

The admixture model of the STRUCTURE software (Pritchard *et al*., 2000) was used to identify potential hybrids in the field population by modelling cluster assignments for K = 1–5 clusters. We made 10 independent runs for each K to confirm consistency across runs. In all simulations, we performed a burn-in period of 10 000 iterations and 1 000 000 Markov chain Monte Carlo iterations. To determine the most likely value of K, we used the ΔK method of (Evanno *et al*., 2005) implemented in STRUCTURE HARVESTER (Earl & Vonholdt, 2012). We combined the results from the 10 replicate runs into one output using CLUMPP software (Jakobsson & Rosenberg, 2007).

### Laboratory cross-breeding experiment

We kept the 71 juveniles collected in 2015 in individual pots and placed them in an incubator until they reached adulthood, in order to have virgin adults. On the basis of available individuals from the different lineages, we used a subset of 66 adults to form 22 trios based on their assignment to mitochondrial lineages, and which we labelled from letter A to V (Table 1). We formed six types of trios with up to two adults per lineage in order to study reproductive strategies when cross-breeding between mitochondrial lineages. In details (and see Table 1), these trios were composed of: (1) two adults of L2 and one adult of L3, (2) two adults of L3 and one adult of L2, (3) three adults of L2, (4) three adults of L3, (5) two adults of L2 and one adult of L1, (6) two adults of L3 and one adult of L1. The trios composed of adults coming from two distinct lineages were replicated four or five times and the trios composed of adults from the same lineage were replicated twice. We collected the cocoons produced by each trio monthly for four months and kept them in an incubator until hatching. Upon hatching, the juveniles were fixed in ethanol before we performed DNA extraction, as described above.

**Table 1.** For each trio, the assemblage of adults according to their mitochondrial lineage, the number of cocoons and juveniles produced and their hatching rate, plus the number and proportion of juveniles genotyped.

| Cross-breeding type |  |  | Offspring |  |  | Genotyping |  | Summary per cross-breeding type |  |
| --- | --- | --- | --- | --- | --- | --- | --- | --- | --- |
| Trio | description | assemblage | # cocoons | # juveniles | % hatching | # genotyped juveniles | % genotyped juveniles | # cocoons (mean ± sd) | % hatching (mean ± sd) |
| A | L2/L3 majority<br>L2 | 2 L2 + 1 L3 | 58 | 51 | 87.9 | 22 | 43.1 | 52.2 ± 9.26 | 39.6 ± 9.63 |
| B |  |  | 63 | 49 | 77.8 | 21 | 42.9 |  |  |
| C |  |  | 39 | 32 | 82.1 | 22 | 68.8 |  |  |
| D |  |  | 53 | 35 | 66.0 | 21 | 60.0 |  |  |
| E |  |  | 48 | 31 | 64.6 | 22 | 71.0 |  |  |
| F | L2/L3 majority<br>L3 | 2 L3 + 1 L2 | 9 | 6 | 66.7 | 6 | 100.0 | 33.6 ± 17.4 | 24.6 ± 14.2 |
| G |  |  | 45 | 24 | 53.3 | 21 | 87.5 |  |  |
| H |  |  | 26 | 23 | 88.5 | 20 | 87.0 |  |  |
| I |  |  | 34 | 24 | 70.6 | 18 | 75.0 |  |  |
| J |  |  | 54 | 46 | 85.2 | 21 | 45.7 |  |  |
| K | L2 pure | 3 L2 | 28 | 21 | 75.0 | 21 | 100 | 40.0 ± 17.0 | 32 ± 15.6 |
| L |  |  | 52 | 43 | 82.7 | 21 | 48.8 |  |  |
| M | L3 pure | 3 L3 | 25 | 15 | 60.0 | 14 | 93.3 | 27.0 ± 2.83 | 19.5 ± 6.36 |
| N |  |  | 29 | 24 | 82.8 | 20 | 83.3 |  |  |
| O | L1/L2 majority<br>L2 | 1 L1 + 2 L2 | 30 | 20 | 66.7 | 20 | 100.0 | 50.2 ± 15.8 | 30 ± 10.2 |
| P |  |  | 68 | 44 | 64.7 | 24 | 54.5 |  |  |
| Q |  |  | 55 | 30 | 54.5 | 24 | 80.0 |  |  |
| R |  |  | 48 | 26 | 54.2 | 25 | 96.2 |  |  |
| S | L1/L3 majority<br>L3 | 1 L1 + 2 L3 | 70 | 38 | 54.3 | 22 | 57.9 | 58.2 ± 10.8 | 34.2 ± 10.3 |
| T |  |  | 64 | 47 | 73.4 | 23 | 48.9 |  |  |
| U |  |  | 46 | 24 | 52.2 | 21 | 87.5 |  |  |
| V |  |  | 53 | 28 | 52.8 | 23 | 82.1 |  |  |
|  |  |  | 997 | 681 | 68.9 ± 12.5 | 452 | 73.3 ± 20.1 |  |  |

#### Parentage assignment using COI mitochondrial marker

We determined the mitochondrial lineage of each hatched juvenile, that is the lineage of the parent that used its female function during mating, using three methods, each specific to each cross-breeding type. (1) For the trios of pure bred of L2 or L3 (trios K to N), the mitochondrial lineage of the hatched juveniles is the one that characterize the trio, since the adults are of the same mitochondrial lineage. (2) For the hatched juveniles from trios composed of L2/L3 and L1 adults (trios O to V corresponding to a mixed between the two colour morphs), we combined High Resolution Melting (HRM) analysis to DNA Barcoding (Bar-HRM), as described by Baudrin *et al*. (2020). This method was shown to discriminate L1 and L2/L3 lineages but not L2 and L3 (Baudrin *et al*., 2020). In brief, the HRM analysis of the putative parents was carried out in triplicates, at the same time as for the juveniles, using MeltDoctor HRM Master Mix (Applied Biosystems) according to the manufacturer protocol in 10 μL reaction volume and using EwD/EwE primers developed by (Bienert *et al*., 2012). (3) For the hatched juveniles produced from the remaining trios, composed of a mixed of L2 and L3 adults of the green morph, we carried out a PCR screening using COI species-specific primers as described by King *et al*. (2010). The COI-AchL2A-F5 + COI-AchL2A-R3, COI-AchL2B-F3 + COI-AchL2B-R3, COI-AchL3-F2 + COI-AchL3-R2 and COI-Ach L1-F4 and COI-Ach L1-R2 were first used to check the method in triplicates on all putative parents. This method allowed us to assign 100% of the putative parents to their correct COI lineage (previously determined by sequencing). It also revealed that all L2 parents could be amplified using L2B specific primers, making the use of L2A specific primers unnecessary. We amplified the DNA of the hatched juveniles in monoplex using two couples of primers, according to the lineage of their putative parents (L2A and L3, L2A and L1, or L3 and L1). Amplifications were performed in 15 μL, containing 5x Flexi Reaction Buffer, 0.125 mM of each dNTP, 1.5 mM MgCl2, 0.5U of GoTaq^®^ Flexi DNA Polymerase (Promega), 0.2μM of each primer and 1 μl of extracted DNA. PCR cycling conditions were 94°C for 3 min, followed by 35 cycles at 94 °C for 30 s, 57 °C (L1, L2A and L3 primers) or 58°C (for L2B primers) for 45 s, 72 °C for 60 s and a final extension at 72 °C for 10 min. We visualized the results on 2% agarose gels. The method used to determine the COI lineage of each hatched juvenile is available in the supplemental information. Finally, we controlled the assignments of 84 hatched juveniles by COI sequencing (supplemental information), using the method described above.

#### Parentage assignment using microsatellite markers

We genotyped 453 out of the 681 hatched juveniles using four microsatellite markers, such as described above. For the two trios F and M that produced few juveniles, we genotyped all the juveniles from the trio F (N juveniles produced = 6) and 14 juveniles of the trio M (N juveniles produced = 15). For trios that produced at least 20 juveniles, we genotyped on average 21.7 ± 1.6 juveniles per trio. Detailed information on the number of juveniles produced and genotyped in each trio are presented in Table 1. Because genotyping errors are an important source of problems for parentage analysis, we duplicated all the PCR results of the adults that were putative parents in the cross-breeding experiments.

The CERVUS v.3.0.7 software was used (i) to calculate the combined non-exclusion probabilities over loci, for first parent (NE-1P), second parent (NE-2P), parent pair (NE-PP), unrelated individuals (NE-I) or siblings (NE-SI), (ii) to determine the confidence of parentage assignments using a likelihood-based approach to assign parental origin combined with simulation parentage analysis, and (iii) to perform parentage assignment. Likelihood score ratios (LOD) estimate the likelihood that the candidate parent is the true parent divided by the likelihood that the candidate parent is not the true parent. Before proceeding to the parentage assignment, simulations were run in CERVUS to determine the distribution of the critical values of LOD score for 80% and 95% confidence levels. The following simulation parameters for 100,000 offspring were chosen: “candidate parents” 3, “prop. sampled” 1, “prop. loci typed” 0.85, and “prop. loci mistyped” 0.01. Confidence levels obtained from simulations were used for true paternity screening of the offspring.

### Reproductive strategies

To investigate differences in mating strategies between the L2 and L3 lineages, we studied reproduction first among the types of trios and then at the level of each parents. These analyses focused on data from trios A to N. We used linear models to investigate differences in cocoons production, hatching rates and the number of hatched juveniles among trios dominated by either L2 (trios A to E and K and L) or L3 (trios F to J and M and N). At the level of each potential parent (of either L2 or L3 lineage), regardless of the trio in which they were involved, we studied the number of juveniles produced by each earthworm and tested for differences according to the lineage of the adults and the number of mates, using a linear model. We also tested whether these lineages consistently preferred to use a function (male or female) in such cross-breeding. This was modelled as a binomial response with, for each parent, a two-vectors variable with the number of offspring produced using the female function and the number of offspring produced using the male function. Note that these analyses were limited to the subset of juveniles we genotyped (see above).

We followed the same procedure to investigate the reproductive isolation between L2 or L3 and L1 lineages. First, we used linear models to test for differences in cocoons production, hatching rates, and the number of hatched juveniles, between trios including one L1 individual (trios O to V) to trios with no L1 (trios A to N). Then, focusing on trios including a L1 parent (trios O to V), we studied the parentage assignment of the juveniles. We specifically tested for differences in the number of juveniles produced from breeding between adults of L2 or L3, to the number of juveniles produced from cross-breeding between an adult of L2 or L3 and an adult of L1, using a linear model. Last, based on the subset of juveniles produced from the crossing between adults of L2 or L3 and L1, we tested whether these lineages consistently preferred to use a function (male or female) in such cross-breeding using a Pearson’s Chi-square test.

All analyses were performed in R version 4.0.2 (R Core Team, 2020).

## Results

### Genetic structure of the field population

We examined and compared the population genetic structure of the *A. chlorotica* agg individuals sampled in the field using their individual assignment based on the mitochondrial COI and the nuclear microsatellites.

#### Mitochondrial genetic variation

We obtained a total of 154 COI sequences from the 157 juveniles and adults sampled in the same habitat. Among them, we identified 20 haplotypes (Genbank accessions MZ930411-30) of which 9, 5, 4 and 2 haplotypes belonged respectively to the L1, L2, L3 and L4 mitochondrial lineages (Fig. 1). In the field population, the majority of the individuals belonged to the L2 (51%) and the L3 (35.7%) lineages, while 10.2% belonged to the L1 lineage and only 3.2% to the L4 lineage (Fig. 2).

**Figure 1.**
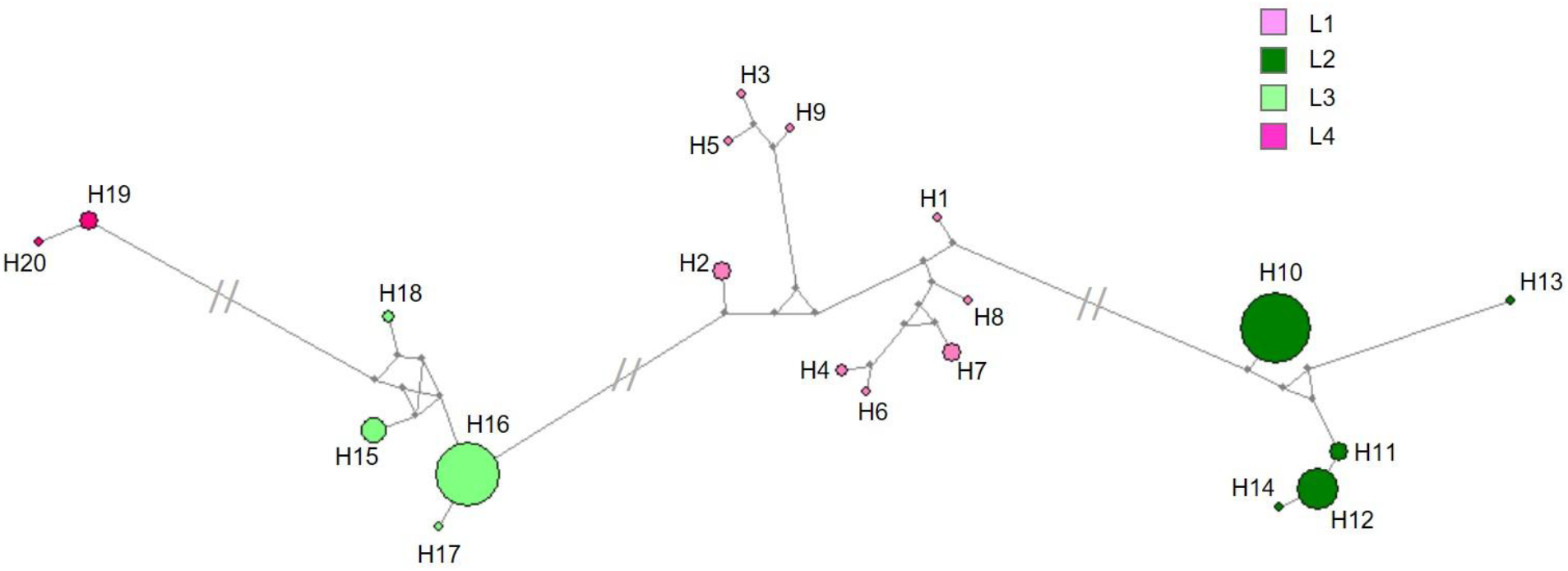
Haplotype network showing the frequency of each COI haplotype belonging to the L1 (H1-H9), L2 (H10-H14), L3 (H15-H18) and L4 (H19-H20) mitochondrial lineages of the *Allolobophora chlorotica* complex in the Preston population, and their relationships. The small grey circles indicate inferred steps not found in the dataset. Connecting lines show mutational pathways between haplotypes. More than 50 mutational steps are indicated.

**Figure 2.**
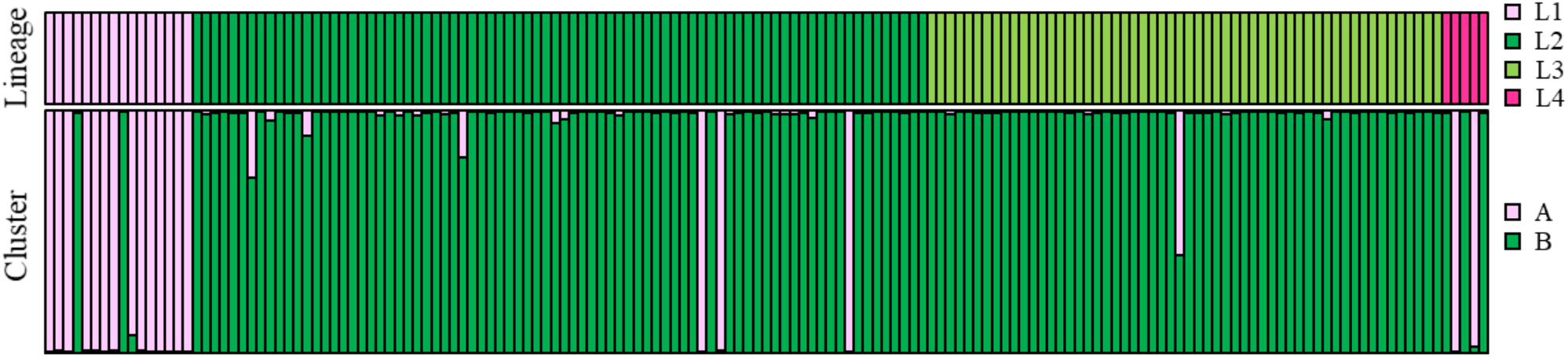
Nuclear clusters and mitochondrial lineages for the field population. Each vertical bar represents one individual. The first row refers to the mitochondrial lineage, the second to the estimated nuclear composition based on the multilocus microsatellite genotype (STRUCTURE software).

#### Nuclear genetic variation

The statistics for the four microsatellite loci used in the study are given in Table 2. Overall polymorphism was high with the number of alleles (*N*_all_) varying from 10 to 25 alleles per locus, *H*_e_ from 0.649 to 0.886, and PIC from 0.605 to 0.873. Null allele frequencies ranged from 0.0306 (Ac476) to 0.1178 (Ac418). The STRUCTURE analysis supported the presence of two nuclear clusters (A and B) within the dataset composed of the 157 individuals sampled in the field (ΔK = 501 for K = 2; Fig. 2). Overall, the nuclear cluster A corresponds to the L1 mitochondrial lineage and the nuclear cluster B to the L2 and L3 mitochondrial lineages, with a few exceptions. Specifically, the case of the L4 mitochondrial lineage was ambiguous since among the five L4 individuals, two were assigned to the nuclear cluster A and three to the nuclear cluster B. This result might be explained by the lower number of microsatellite loci used here, by comparison with Dupont *et al*. (2016; 4 *versus* 5 microsatellite loci). However, this has no consequence on the present study since no L4 individual was used in the cross-breeding experiment. The association was also not categorical for five individuals from the L1, L2 and L3 lineages, whose mitochondrial haplotype did not correspond to their nuclear cluster. Such cases have been previously described as resulting from introgression (Dupont *et al*., 2016). Specifically, two individuals of the L1 mitochondrial lineage were assigned to the nuclear cluster B and three individuals belonging to the L2 mitochondrial lineage were assigned to the nuclear cluster A. Note that these latter three introgressed individuals were used in three different trios (D, E and O, Table 3). Last, three individuals of L2 and one individual of L3 showed admixture above 10%, suggesting recent hybridisation. The L3 hybrid individual which we assigned at 60% to the nuclear cluster B and at 40% to the nuclear cluster A was used in trio B (Table 3).

**Table 2.** Genetic diversity parameters of the four microsatellite loci used in the study and combined non-exclusion probabilities over loci (Combined NE) for first parent (NE-1P), second parent (NE-2P), parent pair (NE-PP), unrelated individual (NE-I) or siblings (NE-SI).

| Locus | $N_{all}$ | $H_o$ | $H_e$ | PIC | NE-1P | NE-2P | NE-PP | NE-I | NE-SI | HW | F(Null) |
| --- | --- | --- | --- | --- | --- | --- | --- | --- | --- | --- | --- |
| Ac127 | 20 | 0.739 | 0.855 | 0.838 | 0.447 | 0.286 | 0.116 | 0.036 | 0.333 | * | 0.0752 |
| Ac170 | 25 | 0.747 | 0.886 | 0.873 | 0.370 | 0.228 | 0.076 | 0.023 | 0.314 | * | 0.0817 |
| Ac418 | 12 | 0.519 | 0.649 | 0.605 | 0.750 | 0.577 | 0.382 | 0.167 | 0.468 | *** | 0.1178 |
| Ac476 | 10 | 0.683 | 0.739 | 0.697 | 0.659 | 0.483 | 0.292 | 0.109 | 0.409 | NS | 0.0306 |
| Mean - Combined NE | 16.75 | 0.672 | 0.782 | 0.753 | 0.08178 | 0.01146 | 0.00099 | 0.00001 | 0.20046 | ND | ND |

\* Significant at the 5% level, \*\*\*significant at the 0.1% level, NS not significant
$N_{all}$ Number of alleles, $H_o$ observed heterozygosity, $H_e$ expected heterozygosity, PIC polymorphic information content, HW exact test of departure from Hardy-Weinberg equilibrium, F(Null) estimated frequency of null alleles, ND Not done.

**Table 3.** Nuclear cluster and parentage assignments. For each trio: the assemblage of adults according to their nuclear cluster such as inferred from the STRUCTURE analysis, the number (No.) of pairs of adults which have produced no juveniles and the mitochondrial lineage and the nuclear cluster of each potential parent, and, the number, lineage and cluster of the adults that produced no juveniles.

| Trio | Nuclear cluster of the adults | Cross without offspring |  |  | Individual without offspring |  |  |
| --- | --- | --- | --- | --- | --- | --- | --- |
|  |  | No. | Lineages | Cluster | No. | Lineage | Cluster |
| A | 3B | 0 | - | - | 0 | - | - |
| B | 2B – 1 hybrid 40%A | 0 | - | - | 0 | - | - |
| C | 3B | 1 | L3-L2 | B-B | 0 | - | - |
| D | 2B – 1 introgressed 100% A | 1 | L2-L2 | B-A† | 0 | - | - |
| E | 2B – 1 introgressed 100% A | 1 | L2-L2 | B-A† | 0 | - | - |
| F | 3B | 2 | L3-L2, L3-L3 | B-B | 1 | L3 | B |
| G | 3B | 0 | - | - | 0 | - | - |
| H | 3B | 1 | L3-L2 | B-B | 0 | - | - |
| I | 3B | 0 | - | - | 0 | - | - |
| J | 3B | 0 | - | - | 0 | - | - |
| K | 3B | 0 | - | - | 0 | - | - |
| L | 3B | 1 | L2-L2 | B-B | 0 | - | - |
| M | 3B | 2 | L3-L3 | B-B | 1 | L3 | B |
| N | 3B | 0 | - | - | 0 | - | - |
| O | 1B - 1 introgressed 100% A – 1A | 1 | L2-L2 | B-A† | 0 | - | - |
| P | 2B – 1A | 1 | L1-L2 | A-B | 0 | - | - |
| Q | 2B – 1A | 2 | L1-L2 | A-B | 1 | L1 | A |
| R | 2B – 1A | 2 | L1-L2 | A-B | 1 | L1 | A |
| S | 2B – 1A | 2 | L1-L3 | A-B | 1 | L1 | A |
| T | 2B – 1A | 0 | - | - | - | - | - |
| U | 2B – 1A | 2 | L1-L3 | A-B | 1 | L1 | A |
| V | 2B – 1A | 2 | L1-L3 | A-B | 1 | L1 | A |

† indicate the introgressed adults.

### Parentage assignment

The simulations in CERVUS resulted in high assignment rates, of 95% and 98% of parental pair assignment, at strict (95%) and relaxed (80%) levels, respectively. Only 2% of simulated offspring remained unassigned. Values of combined non-exclusion probabilities for the first parent, second parent and parent pairs were low, at 0.08178, 0.01815 and 0.00001 respectively. Results per trio are summarized in Table 3 and detailed in the supplemental information. Most of the adults from the trios had offspring, except two L3 individuals in the trios F (L2/L3 majority L3 group) and M (pure L3 group) and five L1 individuals in the trios Q and R (L1/L2 majority L2 group) and S, U and V (L1/L3 majority L3 group). In several trios (A, B, G, I, J, K, N, T) all possible crosses produced offspring, highlighting that individuals frequently mated with the two available partners. It is noteworthy that in the three trios comprising an introgressed individual, one cross in each trio involving this individual did not produce any offspring (Table 3).

### Reproductive strategies

Overall, trios produced 45.32 ± 3.41 cocoons (mean ± se), of which 30.95 ± 2.54 juveniles hatched (mean ± se). The number of cocoons that hatched per trio was proportional to the total number of cocoons it produced (estimate = 0.64 ± 0.08, t = 7.60, p < 0.001, R^2^ = 0.73, Table 1). Based on the parentage assignment of the subset of juveniles genotyped, we noted that 7 adults did not produce any juveniles (Table 3).

To investigate reproductive strategies between adults of the L2 and the L3 lineages, we focused on trios A to N, which involve adults of the L2 and L3 lineages only. We looked at the reproduction among trios and at the individual level of the parent. Among trios, we found that trios with a majority of L2 adults produced significantly more juveniles than the trios with a majority of L3 adults (*F*_1,12_ = 5.42, p = 0.038, Fig. 3). The trios with a majority of L3 produced a significantly lower number of cocoons (estimated difference ± se = −17.00 ± 7.11, t = −2.39, p = 0.034, Fig. 3) but they showed a similar rate of hatching than trios with a majority of L3 (estimated difference ± se = −0.04 ± 0.06, t = −0.69, p = 0.51, Fig. 3). At the individual level of the parent, that is regardless of the trio in which they were involved, we found no difference in the number of juveniles produced between adults of the L2 and L3 lineages (F_1,38_ = 0.35, p = 0.56). However, the adults who mated with both their potential mates produced significantly more juveniles (F_2,38_ = 10.51, p < 0.001). Adults who mated with a unique partner produced 10.43 ± 1.37 juveniles (mean ± se), while adults who mated with both their potential mates produced 15.23 ± 1.70 juveniles (mean ± se). Note that the number of partners did not vary between lineages (Chisq = 3.30, Df = 2, p = 0.19). The parentage assignment of the juveniles also revealed that adults produced significantly more juveniles when mating with adults from the other lineage and this was true for both lineages (for L2 individuals: Chisq = 64.32, Df = 9, p < 0.001, for L3 individuals: Chisq = 43.02, Df = 8, p < 0.001). In such cases of cross-breeding between L2 and L3 lineages, the L2 parent preferentially used their male function while the L3 parent used their female function (Chisq = 6.84, Df = 1, p = 0.009).

**Figure 3.**
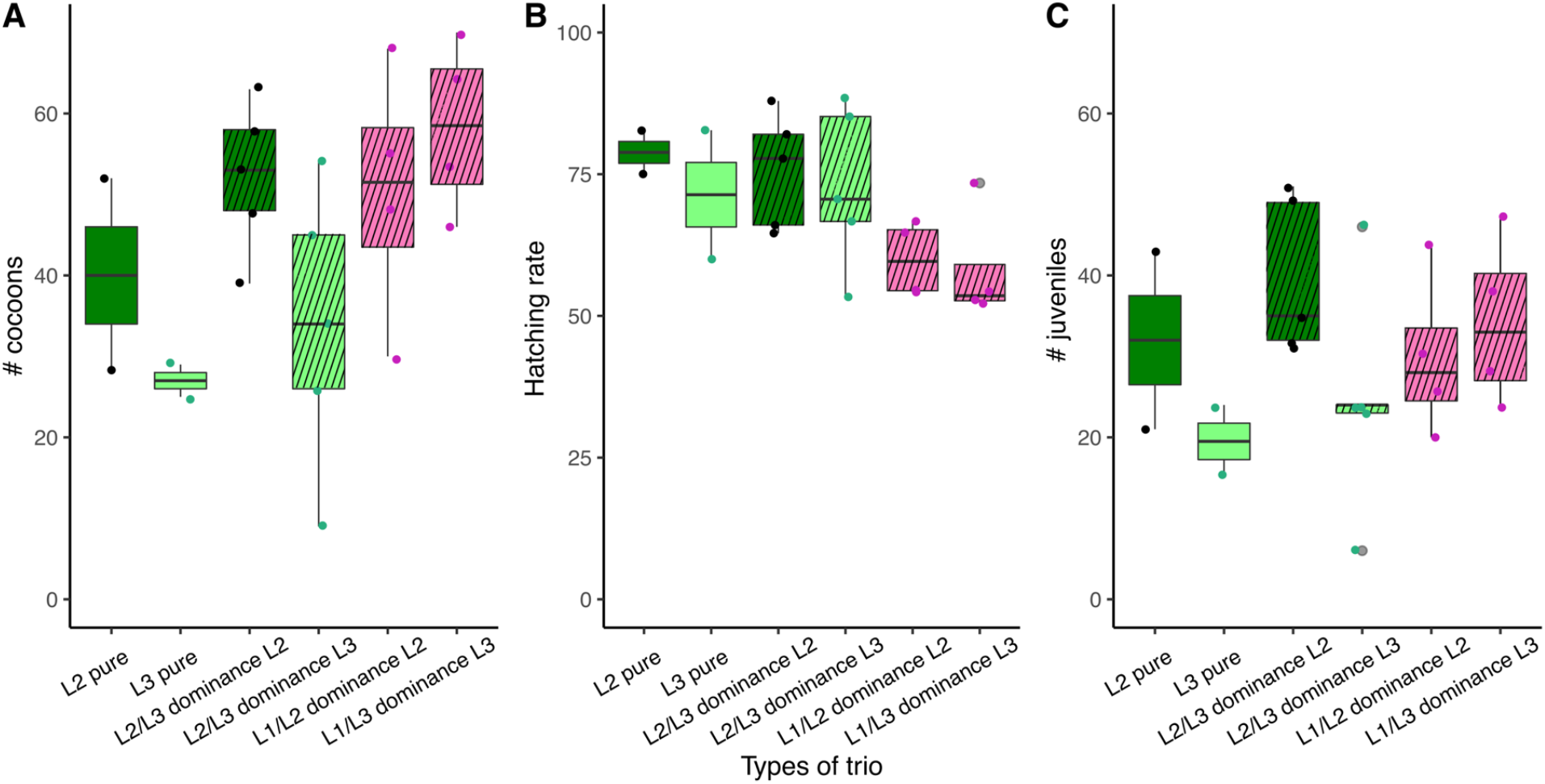
Boxplot of (**A)** the total number of cocoons produced, (**B)** the hatching rate, and (**C)** the number of juveniles, according to each type of trio. In hatched boxes, the values are for the mixed trios of lineages. In light and dark green are the trios of L2 and L3 (of green morph) and in pink are the trios including an adult of L1 (of the pink morph).

When comparing the reproductive rates of trios including one L1 (trios O to V) to trios with no L1 (trios A to N), we observed that trios that involved one L1 adult, produced significantly more cocoons (estimate ± se = 14.04 ± 6.55, t = 2.14, p = 0.04), but their hatching rate was significantly lower (estimate ± se = −0.15 ± 0.045, t = −3.44, p < 0.01), leading to similar numbers of juveniles than in trios without L1 (estimate ± se = 1.84 ± 5.40, t = 0.34, p = 0.74, Fig. 3). The parentage assignment of juveniles in trios including L1 further showed that the cocoons that hatched were mainly from mating between adults of L2 or L3 (at 81.3%, Table 3). The number of juveniles produced by adults of L2 and L3 was significantly higher when they reproduced with an adult of the same lineage than with an L1 adult (estimate ± se = 16.38 ± 2.22, t = 7.39, p < 0.001). It is difficult to draw any solid conclusion about sexual function preferentially used in crossing between L1 and L2 or L3 because only three out of the eight L1 adults produced juveniles from such crossing. Still, based on the available data, 28 of the 34 juveniles were produced using the female function, suggesting that in such crossing L1 adults would preferentially reproduce using their female function (Chisq = 9.90, Df = 2, p = 0.007).

Note that all our results hold, whether or not we considered, for the introgressed individuals, their mitochondrial or nuclear assignment.

## Discussion

The genetic study that we have carried out here, on individuals belonging to different mitochondrial lineages but sampled in the same habitat (i.e. in syntopy) and on their descendants, sheds new light to understand reproductive isolation patterns within the *A. chlorotica* agg. On the one hand, we confirmed that individuals from the two divergent mitochondrial lineages L2 and L3 cannot be distinguished using nuclear marker and mate in the same way as between individuals of the same lineage. On the other hand, we revealed that the sex-function used in the L2-L3 crosses was specific to each lineage. Moreover, we provide additional evidence that mechanisms of post-zygotic reproductive isolation between different but closely related species (L1 and L2/ L3) are at play. Note that our interpretations of the reproductive strategies are for part restricted to the subset of juveniles that have hatched and for which we know the parentage. Still, potential interpretation biases are limited since the juveniles genotyped from each trio were selected at random and that the proportion of genotyped juveniles per trio was relatively high (73.3 ± 20.4 juveniles ranging from 42.9% to 100%). We are, therefore, confident that what we observed is representative of what could have been observed from the genotyping of all individuals.

### Reproductive strategy within the L2/L3 lineages of the *A. chlorotica* agg

The parentage analysis revealed that, over a four-month period, *A. chlorotica* adults can have offspring from multiple partners. Although multiple mating was known in several earthworm species, e.g. in *Eisenia fetida* (Monroy *et al*., 2003), *Lumbricus terrestris* (Butt & Nuutinen, 1998; Michiels *et al*., 2001), and *Hormogaster elisae* (Novo *et al*., 2013), to the best of our knowledge only Novo *et al*. (2013) used similar molecular techniques to examine the fate of sperm after its transfer to a mate. Using microsatellite markers, Novo *et al*. (2013) found cases of multiple paternity in *Hormogaster elisae* and showed that paternity is influenced by the order of copulation. Here, we showed that *A. chlorotica* adults that mated with both their potential mates in the trio produced significantly more juveniles. This result is consistent with the hypothesis of Porto, Velando and Domínguez (2012) who investigated the effect of multiple mating on female reproduction in the earthworm *Eisenia andrei*. These authors found that multiple mating was beneficial for female reproduction, with increased hatching success of the cocoons, and they suggested that this may result from an increase in sperm quantity and/or diversity in the spermathecae of multiple-mated earthworms.

Further, we found that the sex function used in the L2-L3 crosses was specific to each lineage, with the L2 parents that preferentially used their male function and L3 parents their female function. These differences might be the result of a conflict of interest between the mating partners (i.e. the sperm donor and the sperm recipient, Schärer, 2009). It was suggested that sperm donors develop traits that either directly boost the female function of the recipient or disrupt its male function (Schärer, 2009). For instance in the earthworm *Lumbricus terrestris*, the injection of a substance from its setal glands through copulatory setae during mating increases sperm uptake and delays re-mating (Koene *et al*., 2005). By analogy, it could be that during reciprocal copulation and sperm transfer, the L2 individual could transfer a substance that would increase female allocation in the L3 recipient. Understanding more generally these postcopulatory mechanisms of resource allocation between sexual functions and how they may interfere with multiple mating could further our understanding of the sexual selection mechanisms at play in this species. Multiple mating with different partners as well as sperm storage in spermathecae (3 pairs of spermathecae have been described) reported in the morphospecies *A. chlorotica* by Sims and Gerard (1999) further advocate for interference between multiple mating and post-copulatory sexual selection.

### Reproductive isolation between L1 and L2/L3 lineages of the *A. chlorotica* agg

Our results support observations from previous breeding experiments, according to which post-zygotic reproductive isolation processes are occurring between L1 and L2/L3 lineages (Lowe and Butt, 2008). We found that cross-breeding between these lineages produced cocoons. The number of cocoons produced by the trios including a parent of L1 was even higher than by the trios without L1. This is most likely explained by a higher reproduction rate between adults of L2 or L3 and L1 than between adults of L2 and L3, even if these cocoons were, *in fine*, predominantly non-viable. Indeed, we found that the higher production of cocoons in these trios was counterbalanced by a high mortality, as evidenced by their low hatching rate (Table 1). Likewise, Lowe and Butt (2008) found that crosses between adults of the green (L2 and L3) and pink (L1 and L4) morphs were able to reproduce but the viability of their cocoons was substantially reduced, with a hatching rate of 6% and 59% for the green and pink morphs, respectively. It would have been particularly interesting to determine the parents of the cocoons which did not hatch by genotyping. This was attempted, but we were unable to distinguish between the maternal and juvenile DNA in the DNA extracts from the cocoons. Consistently with the observed reproductive isolation, 63% of the L1 adults produced no juveniles and, comparatively, the L2/L3 adults produced significantly more juveniles when reproducing with an adult of the same lineage that with an adult of L1.

Despite evidence of reproductive isolation, we still obtained a few juvenile hybrids of L2/L3 and L1 (in trio O, P and T) from our cross-breeding experiment. Hybrids were also recorded, anecdotally, among individuals from the field population (Fig.2), suggesting that some can reach adulthood. One of these hybrids, with an L3 mitochondrial lineage but assigned at 60% to the cluster B and at 40% to the cluster A, was used in trio B. The parentage analysis revealed that this adult had produced 10 juveniles using only its female function. Sterility of the male function in hybrids has already been proposed from the breeding experiment from Lowe and Butt (2008) and deserves to be further investigated in order to test for Haldane’s rule (i.e. disproportionate hybrid dysfunction in the male function, Haldane, 1922) in this simultaneous hermaphrodite.

Hybrid male sterility would allow an explanation for the existence of introgressed individuals that presented an L2 or L3 mitochondrial lineage but that were assigned to the nuclear cluster A (Fig 2, Table 3, Dupont *et al*., 2016). The case of such introgressed individuals, that we found among those collected in the field population and that were used in three different trios (D, E and O, Table 3), is the most interesting. They were all from the L2 mitochondrial lineage but from the nuclear cluster A. In the trios D and E, composed of L2 and L3 adults with a majority of L2, one of the introgressed individual produced one juvenile using its female function (in trio D), the other produced 7 juveniles using both functions (in trio E: 4 and 3 juveniles produced using, respectively, its male and female function). In trio O, composed of one adult of L1 with two adults of L2, the introgressed individual reproduced using both functions (2 and 4 juveniles produced using, respectively, its male and female function, supplemental information) but only with the L1 adult of the same nuclear cluster (A). Such a small number of juveniles produced from an introgressed parent does not allow any strong conclusion, but still suggests that male function could be restored in introgressed individuals, while it is not active in hybrids. This suggests that, in hybrids, male-specific genes are likely to be non- or mis-expressed while, in introgressed individuals, in which a full nuclear genome of one species has been reconstructed after successive backcrosses, the expression of these genes could be restored.

## Conclusion

It is only recently that works have suggested that earthworms are able to detect the degree of relatedness, the quality and mating status of their partners, and that they are able to fine-tune control of transferred ejaculate volume and cocoon production (Domínguez & Velando, 2013). We believe that further investigations are now needed to interpret these findings in the lights of the sex allocation theory. In simultaneous hermaphrodites such as earthworms, sex allocation is the amount of reproductive effort invested in the male *versus* female function (Charnov, 1996). Basic sex allocation models for simultaneous hermaphrodites generally assume a linear trade-off between the allocation to male and female function, so that higher allocation to one function leads to a proportional decrease in the allocation to the other function (Schärer, 2009). It is expected that individuals show a preference for adopting the sex role that tends to offer the higher potential fitness gain per mating (Anthes *et al*., 2006). The preferred sex role may depend on factors such as the body size, the quality of the partner, the sperm precedence pattern and the mating history of mates (Janicke & Schärer, 2009; Schärer, 2009; van Velzen *et al*., 2009). In the earthworm *E. fetida*, Meyer and Bouwman (1997) showed that within mating pairs there are low cocoon producing individuals that may be considered as sperm donors, and high producers of cocoons that may be considered as sperm receiver. This warrants further investigation, as the difference in sex function used by L2 and L3 adults when cross-breeding could result from such mechanisms of differential sex allocation and may be involved in reproductive isolation in the *A. chlorotica* agg.

## Acknowledgments

The authors are grateful to the Genomic platform of IMRB (Créteil, France) and the PRAMMICS platform of the OSU Efluve (Créteil, France) for providing molecular biology facilities.

