## supplementary material for "Reproductive strategies in a complex of simultaneously hermaphroditic species, the *Allolobophora chlorotica* case study"

### Details of trio composition and parentage assignment in cross-breeding experiment

**Legend :**

The composition of each trio of the cross-breeding experiment (one sheet per trio) is indicated.

The mitochondrial lineage of each individual is showed as well as the method used for this assignment (SEQ = sequencing, PCR = PCR screening using COI species-specific primers, HRM = Bar-HRM method). When the lineage is deduced from that of the parents, it is indicated by "deduced".

When sequencing was carried out, the haplotype of the individual is indicated.

The nuclear cluster of the adults, such as determined by Structure analysis, is indicated.

The parent of each offspring, such as revealed by Cervus analysis, is indicated. When available, the nature of the relationship (mother/father) is specified; otherwise it is indicated NA.

| Trio | Status | Label | MtDNA Lineage | Assignment method | Haplotype | Nuclear Cluster | Parentage assignment | | | |
|---|---|---|---|---|---|---|---|---|---|---|
| A | Adult | Acp02 | L3 | Seq | H15 | B | | | | |
| | | Acp04 | L2 | Seq | H10 | B | | | | |
| | | Acp05 | L2 | Seq | H12 | B | | | | |
| | | | | | | | Mother | Father | Parent 1 | Parent 2 |
| | Offspring | A1 | L2 | PCR+Seq | H10 | NA | Acp04 | Acp05 | NA | NA |
| | | A2 | L2 | PCR+Seq | H10 | NA | Acp04 | Acp05 | NA | NA |
| | | A3 | L2 | PCR+Seq | H10 | NA | Acp04 | Acp05 | NA | NA |
| | | A4 | L2 | PCR+Seq | H10 | NA | Acp04 | Acp05 | NA | NA |
| | | A5 | L2 | PCR+Seq | H10 | NA | Acp04 | Acp05 | NA | NA |
| | | A6 | L2 | PCR+Seq | H10 | NA | Acp04 | Acp05 | NA | NA |
| | | A7 | L2 | PCR+Seq | H10 | NA | Acp04 | Acp05 | NA | NA |
| | | A8 | L3 | PCR+seq | H15 | NA | Acp02 | Acp05 | NA | NA |
| | | A9 | L3 | PCR | NA | NA | Acp02 | Acp05 | NA | NA |
| | | A10 | L3 | Seq+seq | H15 | NA | Acp02 | Acp05 | NA | NA |
| | | A11 | L2 | PCR+Seq | H10 | NA | Acp04 | Acp05 | NA | NA |
| | | A12 | L3 | PCR | NA | NA | Acp02 | Acp05 | NA | NA |
| | | A13 | L3 | PCR | NA | NA | Acp02 | Acp05 | NA | NA |
| | | A14 | L3 | PCR | NA | NA | Acp02 | Acp05 | NA | NA |
| | | A15 | L3 | PCR | NA | NA | Acp02 | Acp05 | NA | NA |
| | | A16 | L2 | Seq | H12 | NA | Acp05 | Acp04 | NA | NA |
| | | A17 | L2 | PCR+Seq | H10 | NA | Acp04 | Acp05 | NA | NA |
| | | A18 | L2 | PCR | NA | NA | Acp04 | Acp02 | NA | NA |
| | | A19 | L3 | PCR | NA | NA | Acp02 | Acp05 | NA | NA |
| | | A20 | L2 | PCR+Seq | H10 | NA | Acp05 | Acp02 | NA | NA |
| | | A21 | L2 | PCR+Seq | H10 | NA | Acp04 | Acp05 | NA | NA |
| | | A22 | L3 | PCR | NA | NA | Acp02 | Acp05 | NA | NA |

| Trio | Status | Label | MtDNA Lineage | Assignment method | Haplotype | Nuclear Cluster | Parentage assignment | | | |
|---|---|---|---|---|---|---|---|---|---|---|
| B | Adult | Acp06 | L2 | Seq | H10 | B | | | | |
| | | Acp07 | L2 | Seq | H11 | B | | | | |
| | | Acp08 | L3 | Seq | H16 | 40% A 60% B | | | | |
| | | | | | | | Mother | Father | Parent 1 | Parent 2 |
| | Offspring | B1 | L3 | PCR+Seq | H16 | NA | Acp08 | Acp07 | NA | NA |
| | | B2 | L2 | PCR+Seq | H11 | NA | Acp07 | Acp06 | NA | NA |
| | | B3 | L3 | PCR | NA | NA | Acp08 | Acp07 | NA | NA |
| | | B4 | L2 | PCR+Seq | H11 | NA | Acp07 | Acp06 | NA | NA |
| | | B5 | L3 | PCR | NA | NA | Acp08 | Acp07 | NA | NA |
| | | B6 | L2 | PCR+Seq | H11 | NA | Acp07 | Acp06 | NA | NA |
| | | B7 | L3 | PCR | NA | NA | Acp08 | Acp06 | NA | NA |
| | | B8 | L2 | PCR+Seq | H10 | NA | Acp06 | Acp07 | NA | NA |
| | | B9 | L2 | PCR+Seq | H10 | NA | Acp06 | Acp07 | NA | NA |
| | | B10 | L2 | PCR+Seq | H10 | NA | Acp06 | Acp07 | NA | NA |
| | | B11 | L2 | PCR+Seq | H10 | NA | Acp06 | Acp07 | NA | NA |
| | | B12 | L3 | PCR | NA | NA | Acp08 | Acp07 | NA | NA |
| | | B13 | L3 | PCR | NA | NA | Acp08 | Acp06 | NA | NA |
| | | B14 | L3 | PCR | NA | NA | Acp08 | Acp07 | NA | NA |
| | | B15 | L2 | PCR+Seq | H11 | NA | Acp07 | Acp06 | NA | NA |
| | | B17 | L2 | PCR+Seq | H11 | NA | Acp07 | Acp06 | NA | NA |
| | | B18 | L2 | PCR+Seq | H10 | NA | Acp06 | Acp07 | NA | NA |
| | | B19 | L3 | PCR | NA | NA | Acp08 | Acp06 | NA | NA |
| | | B20 | L3 | PCR | NA | NA | Acp08 | Acp06 | NA | NA |
| | | B21 | L2 | PCR+Seq | H10 | NA | Acp06 | Acp07 | NA | NA |
| | | B22 | L3 | PCR | NA | NA | Acp08 | Acp07 | NA | NA |

| Trio | Status | Label | MtDNA Lineage | Assignment method | Haplotype | Nuclear Cluster | Parentage assignment | | | |
|---|---|---|---|---|---|---|---|---|---|---|
| C | Adult | Acp09 | L2 | Seq | H10 | B | | | | |
| | | Acp10 | L2 | Seq | H10 | B | | | | |
| | | Acp12 | L3 | Seq | H16 | B | | | | |
| | | | | | | | Mother | Father | Parent 1 | Parent 2 |
| | Offspring | C1 | L2 | PCR+Seq | H10 | NA | Acp10 | Acp12 | NA | NA |
| | | C2 | L2 | PCR | NA | NA | Acp10 | Acp12 | NA | NA |
| | | C3 | L2 | PCR | NA | NA | NA | NA | Acp09 | Acp10 |
| | | C4 | L3 | PCR+Seq | H16 | NA | Acp12 | Acp10 | NA | NA |
| | | C5 | L3 | PCR | NA | NA | Acp12 | Acp10 | NA | NA |
| | | C6 | L2 | PCR | NA | NA | Acp10 | Acp12 | NA | NA |
| | | C7 | L2 | PCR | NA | NA | Acp10 | Acp12 | NA | NA |
| | | C8 | L2 | PCR | NA | NA | NA | NA | Acp09 | Acp10 |
| | | C9 | L2 | Seq | H10 | NA | NA | NA | Acp09 | Acp10 |
| | | C10 | L2 | PCR | NA | NA | Acp10 | Acp12 | NA | NA |
| | | C11 | L3 | PCR | NA | NA | Acp12 | Acp10 | NA | NA |
| | | C12 | L3 | PCR | NA | NA | Acp12 | Acp10 | NA | NA |
| | | C13 | L3 | PCR | NA | NA | Acp12 | Acp10 | NA | NA |
| | | C14 | L3 | PCR | NA | NA | Acp12 | Acp10 | NA | NA |
| | | C15 | L2 | PCR | NA | NA | NA | NA | Acp09 | Acp10 |
| | | C16 | L3 | Seq | H16 | NA | Acp12 | Acp10 | NA | NA |
| | | C17 | L2 | PCR | NA | NA | Acp10 | Acp12 | NA | NA |
| | | C18 | L2 | PCR | NA | NA | NA | NA | Acp09 | Acp10 |
| | | C19 | L2 | PCR | NA | NA | NA | NA | Acp09 | Acp10 |
| | | C20 | L2 | PCR | NA | NA | NA | NA | Acp09 | Acp10 |
| | | C21 | L2 | PCR | NA | NA | Acp12 | Acp10 | NA | NA |
| | | C22 | L3 | PCR | NA | NA | Acp12 | Acp10 | NA | NA |

| Trio | Status | Label | MtDNA Lineage | Assignment method | Haplotype | Nuclear Cluster | Parentage assignment | | | |
|---|---|---|---|---|---|---|---|---|---|---|
| D | Adult | Acp13 | L3 | Seq | H15 | B | | | | |
| | | Acp14 | L2 | Seq | H11 | B | | | | |
| | | Acp15 | L2 | Seq | H10 | A | | | | |
| | | | | | | | Mother | Father | Parent 1 | Parent 2 |
| | Offspring | D1 | L3 | PCR+Seq | H15 | NA | Acp13 | Acp14 | NA | NA |
| | | D2 | L2 | PCR | NA | NA | Acp14 | Acp13 | NA | NA |
| | | D3 | L3 | PCR | NA | NA | Acp13 | Acp14 | NA | NA |
| | | D4 | L3 | PCR | NA | NA | Acp13 | Acp14 | NA | NA |
| | | D5 | L3 | PCR | NA | NA | Acp13 | Acp14 | NA | NA |
| | | D6 | L2 | PCR | NA | NA | Acp14 | Acp13 | NA | NA |
| | | D7 | L2 | PCR | NA | NA | Acp14 | Acp13 | NA | NA |
| | | D8 | L3 | PCR | NA | NA | Acp13 | Acp14 | NA | NA |
| | | D9 | L2 | PCR | NA | NA | Acp14 | Acp13 | NA | NA |
| | | D10 | L3 | PCR | NA | NA | Acp13 | Acp14 | NA | NA |
| | | D12 | L3 | PCR | NA | NA | Acp13 | Acp14 | NA | NA |
| | | D13 | L2 | PCR | NA | NA | Acp14 | Acp13 | NA | NA |
| | | D14 | L2 | PCR | NA | NA | Acp14 | Acp13 | NA | NA |
| | | D15 | L2 | PCR | NA | NA | Acp14 | Acp13 | NA | NA |
| | | D16 | L2 | PCR | NA | NA | Acp14 | Acp13 | NA | NA |
| | | D17 | L2 | PCR | NA | NA | Acp14 | Acp13 | NA | NA |
| | | D18 | L2 | PCR | NA | NA | Acp15 | Acp13 | NA | NA |
| | | D19 | L2 | PCR | NA | NA | Acp14 | Acp13 | NA | NA |
| | | D20 | L3 | PCR | NA | NA | Acp13 | Acp14 | NA | NA |
| | | D21 | L3 | PCR | NA | NA | Acp13 | Acp14 | NA | NA |
| | | D22 | L2 | PCR | NA | NA | Acp14 | Acp13 | NA | NA |

| Trio | Status | Label | MtDNA Lineage | Assignment method | Haplotype | Nuclear Cluster | Parentage assignment | | | |
|---|---|---|---|---|---|---|---|---|---|---|
| E | Adult | Acp17 | L3 | Seq | H16 | B | | | | |
| | | Acp19 | L2 | Seq | H12 | B | | | | |
| | | Acp21 | L2 | Seq | H12 | A | | | | |
| | | | | | | | Mother | Father | Parent 1 | Parent 2 |
| | Offspring | E1 | L2 | PCR+Seq | H12 | NA | Acp19 | Acp17 | NA | NA |
| | | E2 | L2 | PCR | NA | NA | Acp19 | Acp17 | NA | NA |
| | | E3 | L3 | PCR | NA | NA | Acp17 | Acp19 | NA | NA |
| | | E4 | L3 | PCR | NA | NA | Acp17 | Acp21 | NA | NA |
| | | E5 | L2 | PCR | NA | NA | Acp19 | Acp17 | NA | NA |
| | | E6 | L2 | PCR | NA | NA | Acp21 | Acp17 | NA | NA |
| | | E7 | L2 | PCR | NA | NA | Acp19 | Acp17 | NA | NA |
| | | E8 | L2 | PCR | NA | NA | Acp21 | Acp17 | NA | NA |
| | | E9 | L3 | PCR | NA | NA | Acp17 | Acp21 | NA | NA |
| | | E10 | L3 | PCR | NA | NA | Acp17 | Acp21 | NA | NA |
| | | E11 | L2 | PCR | NA | NA | Acp19 | Acp17 | NA | NA |
| | | E12 | L3 | PCR | NA | NA | Acp17 | Acp19 | NA | NA |
| | | E13 | L3 | PCR | NA | NA | Acp17 | Acp19 | NA | NA |
| | | E14 | L3 | PCR | NA | NA | Acp17 | Acp21 | NA | NA |
| | | E15 | L2 | PCR | NA | NA | Acp19 | Acp17 | NA | NA |
| | | E16 | L2 | PCR | NA | NA | Acp19 | Acp17 | NA | NA |
| | | E17 | L3 | PCR | NA | NA | Acp17 | Acp19 | NA | NA |
| | | E18 | L3 | PCR | NA | NA | Acp17 | Acp19 | NA | NA |
| | | E19 | L3 | PCR | NA | NA | Acp17 | Acp19 | NA | NA |
| | | E20 | L2 | PCR | NA | NA | Acp19 | Acp17 | NA | NA |
| | | E21 | L2 | PCR | NA | NA | Acp21 | Acp17 | NA | NA |
| | | E22 | L3 | PCR | NA | NA | Acp17 | Acp19 | NA | NA |

| Trio | Status | Label | MtDNA Lineage | Assignment method | Haplotype | Nuclear Cluster | Parentage assignment | | | |
|---|---|---|---|---|---|---|---|---|---|---|
| F | Adult | Acp18 | L3 | Seq | H16 | B | | | | |
| | | Acp20 | L3 | Seq | H16 | B | | | | |
| | | Acp26 | L2 | Seq | H10 | B | | | | |
| | | | | | | | Mother | Father | Parent 1 | Parent 2 |
| | Offspring | F1 | L3 | PCR | NA | NA | Acp20 | Acp26 | NA | NA |
| | | F2 | L2 | PCR | NA | NA | Acp26 | Acp20 | NA | NA |
| | | F3 | L3 | PCR | NA | NA | Acp20 | Acp26 | NA | NA |
| | | F4 | L2 | PCR | NA | NA | Acp26 | Acp20 | NA | NA |
| | | F5 | L2 | PCR | NA | NA | Acp26 | Acp20 | NA | NA |
| | | F6 | L2 | PCR | NA | NA | Acp26 | Acp20 | NA | NA |

| Trio | Status | Label | MtDNA Lineage | Assignment method | Haplotype | Nuclear Cluster | Parentage assignment | | | |
|---|---|---|---|---|---|---|---|---|---|---|
| G | Adult | Acp22 | L3 | Seq | H16 | B | | | | |
| | | Acp27 | L2 | Seq | H12 | B | | | | |
| | | Acp30 | L3 | Seq | H16 | B | | | | |
| | | | | | | | Mother | Father | Parent 1 | Parent 2 |
| | Offspring | G1 | L2 | PCR+Seq | H12 | NA | Acp27 | Acp22 | NA | NA |
| | | G2 | L2 | PCR | NA | NA | Acp27 | Acp30 | NA | NA |
| | | G3 | L3 | PCR+Seq | H16 | NA | Acp30 | Acp27 | NA | NA |
| | | G4 | L3 | PCR | NA | NA | Acp22 | Acp27 | NA | NA |
| | | G5 | L2 | PCR | NA | NA | Acp27 | Acp30 | NA | NA |
| | | G6 | L3 | PCR | NA | NA | Acp30 | Acp27 | NA | NA |
| | | G7 | L3 | PCR | NA | NA | Acp30 | Acp27 | NA | NA |
| | | G8 | L2 | PCR | NA | NA | Acp27 | Acp30 | NA | NA |
| | | G9 | L3 | PCR | NA | NA | NA | NA | Acp22 | Acp30 |
| | | G10 | L3 | PCR | NA | NA | Acp30 | Acp27 | NA | NA |
| | | G11 | L2 | PCR | NA | NA | Acp27 | Acp30 | NA | NA |
| | | G12 | L2 | PCR | NA | NA | Acp27 | Acp30 | NA | NA |
| | | G13 | L3 | PCR | NA | NA | Acp22 | Acp27 | NA | NA |
| | | G14 | L3 | PCR | NA | NA | Acp22 | Acp27 | NA | NA |
| | | G15 | L2 | PCR | NA | NA | Acp27 | Acp30 | NA | NA |
| | | G17 | L3 | PCR | NA | NA | Acp22 | Acp27 | NA | NA |
| | | G18 | L3 | PCR | NA | NA | Acp22 | Acp27 | NA | NA |
| | | G19 | L3 | PCR | NA | NA | Acp30 | Acp27 | NA | NA |
| | | G20 | L2 | PCR | NA | NA | Acp27 | Acp30 | NA | NA |
| | | G21 | L3 | PCR | NA | NA | Acp22 | Acp27 | NA | NA |
| | | G22 | L3 | PCR | NA | NA | Acp30 | Acp27 | NA | NA |

| Trio | Status | Label | MtDNA Lineage | Assignment method | Haplotype | Nuclear Cluster | Parentage assignment | | | |
|---|---|---|---|---|---|---|---|---|---|---|
| H | Adult | Acp28 | L2 | Seq | H12 | B | | | | |
| | | Acp34 | L3 | Seq | H16 | B | | | | |
| | | Acp36 | L3 | Seq | H16 | B | | | | |
| | | | | | | | Mother | Father | Parent 1 | Parent 2 |
| | Offspring | H1 | L2 | PCR+Seq | H12 | NA | Acp28 | Acp34 | NA | NA |
| | | H2 | L3 | PCR+Seq | H16 | NA | Acp34 | Acp28 | NA | NA |
| | | H3 | L2 | PCR | NA | NA | Acp28 | Acp34 | NA | NA |
| | | H4 | L3 | PCR | NA | NA | Acp34 | Acp28 | NA | NA |
| | | H5 | L2 | PCR | NA | NA | Acp28 | Acp34 | NA | NA |
| | | H6 | L2 | PCR | NA | NA | Acp28 | Acp34 | NA | NA |
| | | H7 | L3 | PCR | NA | NA | Acp34 | Acp28 | NA | NA |
| | | H8 | L2 | PCR | NA | NA | Acp28 | Acp34 | NA | NA |
| | | H9 | L3 | PCR | NA | NA | NA | NA | Acp34 | Acp36 |
| | | H11 | L3 | PCR | NA | NA | Acp34 | Acp28 | NA | NA |
| | | H12 | L3 | PCR | NA | NA | NA | NA | Acp34 | Acp36 |
| | | H13 | L3 | PCR | NA | NA | Acp34 | Acp28 | NA | NA |
| | | H14 | L2 | PCR | NA | NA | Acp28 | Acp34 | NA | NA |
| | | H15 | L2 | PCR | NA | NA | Acp28 | Acp34 | NA | NA |
| | | H16 | L3 | PCR | NA | NA | Acp34 | Acp28 | NA | NA |
| | | H17 | L3 | PCR | NA | NA | Acp34 | Acp28 | NA | NA |
| | | H18 | L2 | PCR | NA | NA | Acp28 | Acp34 | NA | NA |
| | | H19 | L3 | PCR | NA | NA | Acp34 | Acp28 | NA | NA |
| | | H20 | L2 | PCR | NA | NA | Acp28 | Acp34 | NA | NA |
| | | H21 | L3 | PCR | NA | NA | Acp34 | Acp28 | NA | NA |

| Trio | Status | Label | MtDNA Lineage | Assignment method | Haplotype | Nuclear Cluster | Parentage assignment | | | |
|---|---|---|---|---|---|---|---|---|---|---|
| I | Adult | Acp29 | L2 | Seq | H10 | B | | | | |
| | | Acp37 | L3 | Seq | H16 | B | | | | |
| | | Acp38 | L3 | Seq | H16 | B | | | | |
| | | | | | | | Mother | Father | Parent 1 | Parent 2 |
| | Offspring | I1 | L3 | PCR+Seq | H16 | NA | Acp38 | Acp29 | NA | NA |
| | | I2 | L2 | PCR+Seq | H10 | NA | Acp29 | Acp37 | NA | NA |
| | | I3 | L3 | PCR | NA | NA | Acp38 | Acp37 | NA | NA |
| | | I4 | L2 | PCR | NA | NA | Acp29 | Acp37 | NA | NA |
| | | I5 | L3 | PCR | NA | NA | NA | NA | Acp37 | Acp38 |
| | | I6 | L3 | PCR | NA | NA | NA | NA | Acp37 | Acp38 |
| | | I7 | L3 | PCR | NA | NA | NA | NA | Acp37 | Acp38 |
| | | I8 | L3 | PCR | NA | NA | NA | NA | Acp37 | Acp38 |
| | | I9 | L3 | PCR | NA | NA | Acp38 | Acp29 | NA | NA |
| | | I10 | L3 | PCR | NA | NA | NA | NA | Acp37 | Acp38 |
| | | I11 | L3 | PCR | NA | NA | NA | NA | Acp37 | Acp38 |
| | | I12 | L3 | PCR | NA | NA | NA | NA | Acp37 | Acp38 |
| | | I13 | L3 | PCR | NA | NA | NA | NA | Acp37 | Acp38 |
| | | I14 | L3 | PCR | NA | NA | NA | NA | Acp37 | Acp38 |
| | | I15 | L3 | PCR | NA | NA | Acp38 | Acp29 | NA | NA |
| | | I16 | L3 | PCR | NA | NA | Acp38 | Acp29 | NA | NA |
| | | I17 | L3 | PCR | NA | NA | NA | NA | Acp37 | Acp38 |
| | | I18 | L2 | PCR | NA | NA | Acp29 | Acp37 | NA | NA |

| Trio | Status | Label | MtDNA Lineage | Assignment method | Haplotype | Nuclear Cluster | Parentage assignment | | | |
|---|---|---|---|---|---|---|---|---|---|---|
| J | Adult | Acp32 | L2 | Seq | H10 | B | | | | |
| | | Acp39 | L3 | Seq | H16 | B | | | | |
| | | Acp41 | L3 | Seq | H15 | B | | | | |
| | | | | | | | Mother | Father | Parent 1 | Parent 2 |
| | Offspring | J1 | L3 | PCR + seq | H15 | NA | Acp41 | Acp39 | NA | NA |
| | | J2 | L3 | PCR | NA | NA | Acp41 | Acp32 | NA | NA |
| | | J3 | L2 | PCR + seq | H10 | NA | Acp32 | Acp39 | NA | NA |
| | | J4 | L3 | PCR | NA | NA | Acp39 | Acp32 | NA | NA |
| | | J5 | L3 | PCR + seq | H15 | NA | Acp41 | Acp39 | NA | NA |
| | | J6 | L3 | PCR + seq | H16 | NA | Acp39 | Acp41 | NA | NA |
| | | J7 | L3 | PCR + seq | H15 | NA | Acp41 | Acp39 | NA | NA |
| | | J8 | L2 | PCR | NA | NA | Acp32 | Acp39 | NA | NA |
| | | J9 | L3 | PCR | NA | NA | Acp39 | Acp32 | NA | NA |
| | | J10 | L3 | PCR + seq | H15 | NA | Acp41 | Acp39 | NA | NA |
| | | J11 | L3 | PCR + seq | H15 | NA | Acp41 | Acp39 | NA | NA |
| | | J12 | L3 | PCR | NA | NA | Acp41 | Acp32 | NA | NA |
| | | J13 | L3 | PCR | NA | NA | Acp41 | Acp32 | NA | NA |
| | | J14 | L2 | PCR | NA | NA | Acp32 | Acp39 | NA | NA |
| | | J15 | L2 | PCR | NA | NA | Acp32 | Acp41 | NA | NA |
| | | J17 | L3 | PCR + seq | H15 | NA | Acp41 | Acp39 | NA | NA |
| | | J18 | L3 | PCR + seq | H16 | NA | Acp39 | Acp41 | NA | NA |
| | | J19 | L2 | PCR | NA | NA | Acp32 | Acp41 | NA | NA |
| | | J20 | L3 | PCR | NA | NA | Acp41 | Acp32 | NA | NA |
| | | J21 | L3 | PCR + seq | H16 | NA | Acp39 | Acp41 | NA | NA |
| | | J22 | L3 | PCR | NA | NA | Acp39 | Acp32 | NA | NA |

| Trio | Status | Label | MtDNA Lineage | Assignment method | Haplotype | Nuclear Cluster | Parentage assignment | | | |
|---|---|---|---|---|---|---|---|---|---|---|
| K | Adult | Acp33 | L2 | Seq | H10 | B | | | | |
| | | Acp40 | L2 | Seq | H10 | B | | | | |
| | | Acp44 | L2 | Seq | H10 | B | | | | |
| | | | | | | | Mother | Father | Parent 1 | Parent 2 |
| | Offspring | K1 | L2 | deduced | NA | NA | NA | NA | Acp33 | Acp44 |
| | | K2 | L2 | deduced | NA | NA | NA | NA | Acp33 | Acp40 |
| | | K3 | L2 | deduced | NA | NA | NA | NA | Acp33 | Acp44 |
| | | K4 | L2 | deduced | NA | NA | NA | NA | Acp40 | Acp44 |
| | | K5 | L2 | deduced | NA | NA | NA | NA | Acp33 | Acp44 |
| | | K6 | L2 | deduced | NA | NA | NA | NA | Acp40 | Acp44 |
| | | K7 | L2 | deduced | NA | NA | NA | NA | Acp40 | Acp44 |
| | | K8 | L2 | deduced | NA | NA | NA | NA | Acp33 | Acp44 |
| | | K9 | L2 | deduced | NA | NA | NA | NA | Acp33 | Acp44 |
| | | K10 | L2 | deduced | NA | NA | NA | NA | Acp33 | Acp44 |
| | | K11 | L2 | deduced | NA | NA | NA | NA | Acp33 | Acp44 |
| | | K12 | L2 | deduced | NA | NA | NA | NA | Acp40 | Acp44 |
| | | K13 | L2 | deduced | NA | NA | NA | NA | Acp33 | Acp44 |
| | | K14 | L2 | deduced | NA | NA | NA | NA | Acp40 | Acp44 |
| | | K15 | L2 | deduced | NA | NA | NA | NA | Acp40 | Acp44 |
| | | K16 | L2 | deduced | NA | NA | NA | NA | Acp40 | Acp44 |
| | | K17 | L2 | deduced | NA | NA | NA | NA | Acp33 | Acp44 |
| | | K18 | L2 | deduced | NA | NA | NA | NA | Acp40 | Acp44 |
| | | K19 | L2 | deduced | NA | NA | NA | NA | Acp40 | Acp44 |
| | | K20 | L2 | deduced | NA | NA | NA | NA | Acp40 | Acp44 |
| | | K21 | L2 | deduced | NA | NA | NA | NA | Acp40 | Acp44 |
| | | K22 | L2 | deduced | NA | NA | NA | NA | Acp40 | Acp44 |

| Trio | Status | Label | MtDNA Lineage | Assignment method | Haplotype | Nuclear Cluster | Parentage assignment | | | |
|---|---|---|---|---|---|---|---|---|---|---|
| L | Adult | Acp45 | L2 | Seq | H10 | B | | | | |
| | | Acp46 | L2 | Seq | H10 | B | | | | |
| | | Acp47 | L2 | Seq | H10 | B | | | | |
| | | | | | | | Mother | Father | Parent 1 | Parent 2 |
| | Offspring | L1 | L2 | deduced | NA | NA | NA | NA | Acp45 | Acp46 |
| | | L2 | L2 | deduced | NA | NA | NA | NA | Acp45 | Acp46 |
| | | L3 | L2 | deduced | NA | NA | NA | NA | Acp45 | Acp46 |
| | | L4 | L2 | deduced | NA | NA | NA | NA | Acp45 | Acp46 |
| | | L5 | L2 | deduced | NA | NA | NA | NA | Acp45 | Acp47 |
| | | L6 | L2 | deduced | NA | NA | NA | NA | Acp45 | Acp46 |
| | | L7 | L2 | deduced | NA | NA | NA | NA | Acp45 | Acp46 |
| | | L8 | L2 | deduced | NA | NA | NA | NA | Acp45 | Acp46 |
| | | L9 | L2 | deduced | NA | NA | NA | NA | Acp45 | Acp47 |
| | | L10 | L2 | deduced | NA | NA | NA | NA | Acp45 | Acp47 |
| | | L11 | L2 | deduced | NA | NA | NA | NA | Acp45 | Acp46 |
| | | L12 | L2 | deduced | NA | NA | NA | NA | Acp45 | Acp46 |
| | | L13 | L2 | deduced | NA | NA | NA | NA | Acp45 | Acp47 |
| | | L14 | L2 | deduced | NA | NA | NA | NA | Acp45 | Acp47 |
| | | L15 | L2 | deduced | NA | NA | NA | NA | Acp45 | Acp46 |
| | | L17 | L2 | deduced | NA | NA | NA | NA | Acp45 | Acp47 |
| | | L18 | L2 | deduced | NA | NA | NA | NA | Acp45 | Acp47 |
| | | L19 | L2 | deduced | NA | NA | NA | NA | Acp45 | Acp46 |
| | | L20 | L2 | deduced | NA | NA | NA | NA | Acp45 | Acp47 |
| | | L21 | L2 | deduced | NA | NA | NA | NA | Acp45 | Acp46 |
| | | L22 | L2 | deduced | NA | NA | NA | NA | Acp45 | Acp46 |

| Trio | Status | Label | MtDNA Lineage | Assignment method | Haplotype | Nuclear Cluster | Parentage assignment | | | |
|---|---|---|---|---|---|---|---|---|---|---|
| M | Adult | **Acp42** | L3 | Seq | H15 | B | | | | |
| | | **Acp43** | L3 | Seq | H16 | B | | | | |
| | | **Acp49** | L3 | Seq | H16 | B | | | | |
| | | | | | | | Mother | Father | Parent 1 | Parent 2 |
| | Offspring | **M1** | L3 | deduced | NA | NA | NA | NA | Acp42 | Acp43 |
| | | **M2** | L3 | deduced | NA | NA | NA | NA | Acp42 | Acp43 |
| | | **M3** | L3 | deduced | NA | NA | NA | NA | Acp42 | Acp43 |
| | | **M4** | L3 | deduced | NA | NA | NA | NA | Acp42 | Acp43 |
| | | **M5** | L3 | deduced | NA | NA | NA | NA | Acp42 | Acp43 |
| | | **M6** | L3 | deduced | NA | NA | NA | NA | Acp42 | Acp43 |
| | | **M7** | L3 | deduced | NA | NA | NA | NA | Acp42 | Acp43 |
| | | **M8** | L3 | deduced | NA | NA | NA | NA | Acp42 | Acp43 |
| | | **M9** | L3 | deduced | NA | NA | NA | NA | Acp42 | Acp43 |
| | | **M10** | L3 | deduced | NA | NA | NA | NA | Acp42 | Acp43 |
| | | **M12** | L3 | deduced | NA | NA | NA | NA | Acp42 | Acp43 |
| | | **M13** | L3 | deduced | NA | NA | NA | NA | Acp42 | Acp43 |
| | | **M14** | L3 | deduced | NA | NA | NA | NA | Acp42 | Acp43 |
| | | **M16** | L3 | deduced | NA | NA | NA | NA | Acp42 | Acp43 |

| Trio | Status | Label | MtDNA Lineage | Assignment method | Haplotype | Nuclear Cluster | Parentage assignment | | | |
|---|---|---|---|---|---|---|---|---|---|---|
| N | Adult | Acp50 | L3 | Seq | H15 | B | | | | |
| | | Acp52 | L3 | Seq | H16 | B | | | | |
| | | Acp54 | L3 | Seq | H16 | B | | | | |
| | | | | | | | Mother | Father | Parent 1 | Parent 2 |
| | Offspring | N2 | L3 | deduced | NA | NA | NA | NA | Acp50 | Acp52 |
| | | N3 | L3 | deduced | NA | NA | NA | NA | Acp50 | Acp52 |
| | | N4 | L3 | deduced | NA | NA | NA | NA | Acp50 | Acp54 |
| | | N5 | L3 | deduced | NA | NA | NA | NA | Acp50 | Acp54 |
| | | N6 | L3 | deduced | NA | NA | NA | NA | Acp50 | Acp54 |
| | | N7 | L3 | deduced | NA | NA | NA | NA | Acp52 | Acp54 |
| | | N8 | L3 | deduced | NA | NA | NA | NA | Acp50 | Acp54 |
| | | N9 | L3 | deduced | NA | NA | NA | NA | Acp50 | Acp54 |
| | | N10 | L3 | deduced | NA | NA | NA | NA | Acp50 | Acp54 |
| | | N11 | L3 | deduced | NA | NA | NA | NA | Acp50 | Acp54 |
| | | N12 | L3 | deduced | NA | NA | NA | NA | Acp50 | Acp54 |
| | | N13 | L3 | deduced | NA | NA | NA | NA | Acp50 | Acp52 |
| | | N14 | L3 | deduced | NA | NA | NA | NA | Acp52 | Acp54 |
| | | N15 | L3 | deduced | NA | NA | NA | NA | Acp50 | Acp52 |
| | | N17 | L3 | deduced | NA | NA | NA | NA | Acp50 | Acp52 |
| | | N18 | L3 | deduced | NA | NA | NA | NA | Acp50 | Acp54 |
| | | N19 | L3 | deduced | NA | NA | NA | NA | Acp50 | Acp52 |
| | | N20 | L3 | deduced | NA | NA | NA | NA | Acp50 | Acp54 |
| | | N21 | L3 | deduced | NA | NA | NA | NA | Acp50 | Acp52 |
| | | N22 | L3 | deduced | NA | NA | NA | NA | Acp50 | Acp54 |

| Trio | Status | Label | MtDNA Lineage | Assignment method | Haplotype | Nuclear Cluster | Parentage assignment | | | |
|---|---|---|---|---|---|---|---|---|---|---|
| O | Adult | Acp 03 | L1 | Seq | H1 | A | | | | |
| | | Acp 48 | L2 | Seq | H12 | A | | | | |
| | | Acp 59 | L2 | Seq | H10 | B | | | | |
| | | | | | | | Mother | Father | Parent 1 | Parent 2 |
| | Offspring | O1 | L1 | HRM | NA | NA | Acp 03 | Acp 59 | NA | NA |
| | | O3 | L1 | HRM | NA | NA | Acp 03 | Acp 59 | NA | NA |
| | | O4 | L1 | HRM | NA | NA | Acp 03 | Acp 59 | NA | NA |
| | | O5 | L2 | Seq | H12 | NA | Acp 48 | Acp 03 | NA | NA |
| | | O6 | L2 | HRM | NA | NA | Acp 48 | Acp 03 | NA | NA |
| | | O7 | L1 | HRM +Seq | H1 | NA | Acp 03 | Acp 59 | NA | NA |
| | | O8 | L2 | HRM | NA | NA | Acp 03 | Acp 48 | NA | NA |
| | | O9 | L2 | HRM | NA | NA | Acp 03 | Acp 48 | NA | NA |
| | | O10 | L1 | HRM | NA | NA | Acp 03 | Acp 59 | NA | NA |
| | | O11 | L1 | HRM | NA | NA | Acp 03 | Acp 59 | NA | NA |
| | | O12 | L2 | HRM | NA | NA | Acp 48 | Acp 03 | NA | NA |
| | | O13 | L2 | HRM | NA | NA | Acp 59 | Acp 03 | NA | NA |
| | | O14 | L1 | HRM | NA | NA | Acp 03 | Acp 59 | NA | NA |
| | | O15 | L1 | HRM | NA | NA | Acp 03 | Acp 59 | NA | NA |
| | | O16 | L1 | HRM | NA | NA | Acp 03 | Acp 59 | NA | NA |
| | | O17 | L1 | HRM | NA | NA | Acp 03 | Acp 59 | NA | NA |
| | | O18 | L1 | HRM | NA | NA | Acp 03 | Acp 59 | NA | NA |
| | | O19 | L2 | PCR | NA | NA | Acp 48 | Acp 03 | NA | NA |
| | | O20 | L2 | PCR | NA | NA | Acp 59 | Acp 03 | NA | NA |
| | | O21 | L1 | PCR | NA | NA | Acp 03 | Acp 59 | NA | NA |

| Trio | Status | Label | MtDNA Lineage | Assignment method | Haplotype | Nuclear Cluster | Parentage assignment | | | |
|---|---|---|---|---|---|---|---|---|---|---|
| P | Adult | Acp61 | L2 | Seq | H10 | B | | | | |
| | | Acp63 | L2 | Seq | H10 | B | | | | |
| | | Acp11 | L1 | seq | H2 | A | | | | |
| | | | | | | | Mother | Father | Parent 1 | Parent 2 |
| | Offspring | P1 | L2 | HRM | | | NA | NA | Acp61 | Acp63 |
| | | P2 | L2 | HRM | | | NA | NA | Acp61 | Acp63 |
| | | P3 | L2 | HRM | | | NA | NA | Acp61 | Acp63 |
| | | P4 | L2 | HRM | | | NA | NA | Acp61 | Acp63 |
| | | P5 | L1 | HRM + Seq | H2 | | Acp 11 | Acp61 | NA | NA |
| | | P6 | L2 | HRM | | | NA | NA | Acp61 | Acp63 |
| | | P7 | L2 | HRM | | | NA | NA | Acp61 | Acp63 |
| | | P8 | L1 | HRM + Seq | H2 | | Acp 11 | Acp61 | NA | NA |
| | | P9 | L1 | HRM | | | Acp 11 | Acp61 | NA | NA |
| | | P10 | L2 | HRM | | | NA | NA | Acp61 | Acp63 |
| | | P11 | L2 | HRM | | | NA | NA | Acp61 | Acp63 |
| | | P12 | L2 | HRM | | | NA | NA | Acp61 | Acp63 |
| | | P13 | L2 | HRM | | | NA | NA | Acp61 | Acp63 |
| | | P14 | L1 | HRM | | | Acp 11 | Acp61 | | |
| | | P15 | L2 | HRM | | | NA | NA | Acp61 | Acp63 |
| | | P16 | L2 | HRM | | | NA | NA | Acp61 | Acp63 |
| | | P17 | L2 | HRM | | | NA | NA | Acp61 | Acp63 |
| | | P18 | L2 | HRM | | | NA | NA | Acp61 | Acp63 |
| | | P19 | L2 | PCR | | | NA | NA | Acp61 | Acp63 |
| | | P20 | L2 | PCR | | | NA | NA | Acp61 | Acp63 |
| | | P21 | L1 | seq | H2 | | Acp 11 | Acp61 | NA | NA |
| | | P22 | L1 | seq | H2 | | Acp 11 | Acp61 | NA | NA |
| | | P23 | L2 | PCR | | | NA | NA | Acp61 | Acp63 |
| | | P24 | L2 | PCR | | | NA | NA | Acp61 | Acp63 |

| Trio | Status | Label | MtDNA Lineage | Assignment method | Haplotype | Nuclear Cluster | Parentage assignment | | | |
|---|---|---|---|---|---|---|---|---|---|---|
| Q | Adult | **Acp65** | L2 | Seq | H12 | B | | | | |
| | | **Acp67** | L2 | Seq | H12 | B | | | | |
| | | **Acp23** | L1 | Seq | H3 | A | | | | |
| | | | | | | | **Mother** | **Father** | **Parent 1** | **Parent 2** |
| | Offspring | **Q1** | L2 | HRM + Seq | H12 | NA | NA | NA | Acp65 | Acp67 |
| | | **Q2** | L2 | HRM | NA | NA | NA | NA | Acp65 | Acp67 |
| | | **Q3** | L2 | HRM | NA | NA | NA | NA | Acp65 | Acp67 |
| | | **Q4** | L2 | HRM | NA | NA | NA | NA | Acp65 | Acp67 |
| | | **Q5** | L2 | HRM | NA | NA | NA | NA | Acp65 | Acp67 |
| | | **Q6** | L2 | HRM | NA | NA | NA | NA | Acp65 | Acp67 |
| | | **Q7** | L2 | HRM | NA | NA | NA | NA | Acp65 | Acp67 |
| | | **Q8** | L2 | HRM | NA | NA | NA | NA | Acp65 | Acp67 |
| | | **Q9** | L2 | HRM | NA | NA | NA | NA | Acp65 | Acp67 |
| | | **Q10** | L2 | HRM | NA | NA | NA | NA | Acp65 | Acp67 |
| | | **Q11** | L2 | HRM | NA | NA | NA | NA | Acp65 | Acp67 |
| | | **Q12** | L2 | HRM | NA | NA | NA | NA | Acp65 | Acp67 |
| | | **Q13** | L2 | HRM | NA | NA | NA | NA | Acp65 | Acp67 |
| | | **Q14** | L2 | HRM | NA | NA | NA | NA | Acp65 | Acp67 |
| | | **Q15** | L2 | HRM | NA | NA | NA | NA | Acp65 | Acp67 |
| | | **Q16** | L2 | HRM | NA | NA | NA | NA | Acp65 | Acp67 |
| | | **Q17** | L2 | HRM | NA | NA | NA | NA | Acp65 | Acp67 |
| | | **Q18** | L2 | HRM | NA | NA | NA | NA | Acp65 | Acp67 |
| | | **Q19** | L2 | PCR | NA | NA | NA | NA | Acp65 | Acp67 |
| | | **Q20** | L2 | PCR | NA | NA | NA | NA | Acp65 | Acp67 |
| | | **Q21** | L2 | PCR | NA | NA | NA | NA | Acp65 | Acp67 |
| | | **Q22** | L2 | PCR | NA | NA | NA | NA | Acp65 | Acp67 |
| | | **Q23** | L2 | PCR | NA | NA | NA | NA | Acp65 | Acp67 |
| | | **Q24** | L2 | PCR | NA | NA | NA | NA | Acp65 | Acp67 |

| Trio | Status | Label | MtDNA Lineage | Assignment method | Haplotype | Nuclear Cluster | Parentage assignment | | | |
|---|---|---|---|---|---|---|---|---|---|---|
| R | Adult | Acp68 | L2 | Seq | H10 | B | | | | |
| | | Acp69 | L2 | Seq | H10 | B | | | | |
| | | Acp35 | L1 | Seq | H4 | A | | | | |
| | | | | | | | **Mother** | **Father** | **Parent 1** | **Parent 2** |
| | Offspring | R1 | L2 | HRM + Seq | H10 | NA | NA | NA | Acp68 | Acp69 |
| | | R2 | L2 | HRM | NA | NA | NA | NA | Acp68 | Acp69 |
| | | R3 | L2 | HRM | NA | NA | NA | NA | Acp68 | Acp69 |
| | | R4 | L2 | HRM | NA | NA | NA | NA | Acp68 | Acp69 |
| | | R5 | L2 | HRM | NA | NA | NA | NA | Acp68 | Acp69 |
| | | R6 | L2 | HRM | NA | NA | NA | NA | Acp68 | Acp69 |
| | | R7 | L2 | HRM | NA | NA | NA | NA | Acp68 | Acp69 |
| | | R8 | L2 | HRM | NA | NA | NA | NA | Acp68 | Acp69 |
| | | R9 | L2 | HRM | NA | NA | NA | NA | Acp68 | Acp69 |
| | | R10 | L2 | HRM | NA | NA | NA | NA | Acp68 | Acp69 |
| | | R11 | L2 | HRM | NA | NA | NA | NA | Acp68 | Acp69 |
| | | R12 | L2 | HRM | NA | NA | NA | NA | Acp68 | Acp69 |
| | | R13 | L2 | HRM | NA | NA | NA | NA | Acp68 | Acp69 |
| | | R14 | L2 | HRM | NA | NA | NA | NA | Acp68 | Acp69 |
| | | R15 | L2 | HRM | NA | NA | NA | NA | Acp68 | Acp69 |
| | | R16 | L2 | HRM | NA | NA | NA | NA | Acp68 | Acp69 |
| | | R17 | L2 | HRM | NA | NA | NA | NA | Acp68 | Acp69 |
| | | R18 | L2 | HRM | NA | NA | NA | NA | Acp68 | Acp69 |
| | | R19 | L2 | PCR | NA | NA | NA | NA | Acp68 | Acp69 |
| | | R20 | L2 | PCR | NA | NA | NA | NA | Acp68 | Acp69 |
| | | R21 | L2 | PCR | NA | NA | NA | NA | Acp68 | Acp69 |
| | | R22 | L2 | PCR | NA | NA | NA | NA | Acp68 | Acp69 |
| | | R23 | L2 | PCR | NA | NA | NA | NA | Acp68 | Acp69 |
| | | R24 | L2 | PCR | NA | NA | NA | NA | Acp68 | Acp69 |
| | | R25 | L2 | PCR | NA | NA | NA | NA | Acp68 | Acp69 |

| Trio | Status | Label | MtDNA Lineage | Assignment method | Haplotype | Nuclear Cluster | Parentage assignment | | | |
|---|---|---|---|---|---|---|---|---|---|---|
| S | Adult | Acp55 | L3 | Seq | H16 | B | | | | |
| | | Acp57 | L3 | Seq | H16 | B | | | | |
| | | Acp51 | L1 | Seq | H5 | A | | | | |
| | | | | | | | Mother | Father | Parent 1 | Parent 2 |
| | Offspring | S1 | L3 | HRM + Seq | H16 | NA | NA | NA | Acp55 | Acp57 |
| | | S3 | L3 | HRM | NA | NA | NA | NA | Acp55 | Acp57 |
| | | S4 | L3 | HRM | NA | NA | NA | NA | Acp55 | Acp57 |
| | | S5 | L3 | HRM | NA | NA | NA | NA | Acp55 | Acp57 |
| | | S6 | L3 | HRM | NA | NA | NA | NA | Acp55 | Acp57 |
| | | S7 | L3 | HRM | NA | NA | NA | NA | Acp55 | Acp57 |
| | | S8 | L3 | HRM | NA | NA | NA | NA | Acp55 | Acp57 |
| | | S9 | L3 | HRM | NA | NA | NA | NA | Acp55 | Acp57 |
| | | S10 | L3 | HRM | NA | NA | NA | NA | Acp55 | Acp57 |
| | | S11 | L3 | HRM | NA | NA | NA | NA | Acp55 | Acp57 |
| | | S12 | L3 | HRM | NA | NA | NA | NA | Acp55 | Acp57 |
| | | S13 | L3 | HRM | NA | NA | NA | NA | Acp55 | Acp57 |
| | | S14 | L3 | HRM | NA | NA | NA | NA | Acp55 | Acp57 |
| | | S15 | L3 | HRM | NA | NA | NA | NA | Acp55 | Acp57 |
| | | S16 | L3 | HRM | NA | NA | NA | NA | Acp55 | Acp57 |
| | | S17 | L3 | HRM | NA | NA | NA | NA | Acp55 | Acp57 |
| | | S18 | L3 | HRM | NA | NA | NA | NA | Acp55 | Acp57 |
| | | S19 | L3 | PCR | NA | NA | NA | NA | Acp55 | Acp57 |
| | | S20 | L3 | PCR | NA | NA | NA | NA | Acp55 | Acp57 |
| | | S21 | L3 | PCR | NA | NA | NA | NA | Acp55 | Acp57 |
| | | S22 | L3 | PCR | NA | NA | NA | NA | Acp55 | Acp57 |
| | | S23 | L3 | PCR | NA | NA | NA | NA | Acp55 | Acp57 |

| Trio | Status | Label | MtDNA Lineage | Assignment method | Haplotype | Nuclear Cluster | Parentage assignment | | | |
|---|---|---|---|---|---|---|---|---|---|---|
| T | Adult | Acp58 | L3 | Seq | H16 | B | | | | |
| | | Acp60 | L3 | Seq | H16 | B | | | | |
| | | Acp53 | L1 | Seq | H6 | A | | | | |
| | | | | | | | Mother | Father | Parent 1 | Parent 2 |
| | Offspring | T1 | L3 | HRM + Seq | H16 | NA | NA | NA | Acp58 | Acp60 |
| | | T2 | L3 | HRM | NA | NA | NA | NA | Acp58 | Acp60 |
| | | T3 | L3 | HRM | NA | NA | NA | NA | Acp58 | Acp60 |
| | | T4 | L3 | HRM | NA | NA | NA | NA | Acp58 | Acp60 |
| | | T5 | L1 | HRM + Seq | H6 | NA | Acp 53 | Acp60 | NA | NA |
| | | T6 | L1 | HRM | NA | NA | Acp 53 | Acp60 | NA | NA |
| | | T7 | L3 | HRM | NA | NA | NA | NA | Acp58 | Acp60 |
| | | T8 | L3 | HRM | NA | NA | NA | NA | Acp58 | Acp60 |
| | | T9 | L3 | HRM | NA | NA | NA | NA | Acp58 | Acp60 |
| | | T10 | L1 | HRM + Seq | H6 | NA | Acp 53 | Acp60 | NA | NA |
| | | T11 | L3 | HRM | NA | NA | NA | NA | Acp58 | Acp60 |
| | | T12 | L3 | HRM | NA | NA | NA | NA | Acp58 | Acp60 |
| | | T13 | L1 | HRM | NA | NA | Acp 53 | Acp60 | NA | NA |
| | | T14 | L1 | HRM | NA | NA | Acp 53 | Acp60 | NA | NA |
| | | T15 | L3 | HRM | NA | NA | NA | NA | Acp58 | Acp60 |
| | | T16 | L3 | HRM | NA | NA | NA | NA | Acp58 | Acp60 |
| | | T17 | L1 | HRM | NA | NA | Acp 53 | Acp58 | NA | NA |
| | | T18 | L3 | HRM | NA | NA | NA | NA | Acp58 | Acp60 |
| | | T19 | L1 | PCR | NA | NA | Acp53 | Acp60 | NA | NA |
| | | T20 | L1 | PCR | NA | NA | Acp53 | Acp60 | NA | NA |
| | | T21 | L3 | PCR | NA | NA | NA | NA | Acp58 | Acp60 |
| | | T22 | L3 | PCR | NA | NA | NA | NA | Acp58 | Acp60 |
| | | T23 | L3 | PCR | NA | NA | NA | NA | Acp58 | Acp60 |

| Trio | Status | Label | MtDNA Lineage | Assignment method | Haplotype | Nuclear Cluster | Parentage assignment | | | |
|---|---|---|---|---|---|---|---|---|---|---|
| U | Adult | Acp62 | L3 | Seq | H16 | B | | | | |
| | | Acp64 | L3 | Seq | H16 | B | | | | |
| | | Acp56 | L1 | seq | H2 | A | | | | |
| | | | | | | | Mother | Father | Parent 1 | Parent 2 |
| | Offspring | U1 | L3 | HRM+Seq | H16 | NA | NA | NA | Acp 62 | Acp 64 |
| | | U2 | L3 | HRM | NA | NA | NA | NA | Acp 62 | Acp 64 |
| | | U3 | L3 | HRM | NA | NA | NA | NA | Acp 62 | Acp 64 |
| | | U4 | L3 | HRM | NA | NA | NA | NA | Acp 62 | Acp 64 |
| | | U5 | L3 | HRM | NA | NA | NA | NA | Acp 62 | Acp 64 |
| | | U6 | L3 | HRM | NA | NA | NA | NA | Acp 62 | Acp 64 |
| | | U7 | L3 | HRM | NA | NA | NA | NA | Acp 62 | Acp 64 |
| | | U8 | L3 | HRM | NA | NA | NA | NA | Acp 62 | Acp 64 |
| | | U9 | L3 | HRM | NA | NA | NA | NA | Acp 62 | Acp 64 |
| | | U10 | L3 | HRM | NA | NA | NA | NA | Acp 62 | Acp 64 |
| | | U11 | L3 | HRM | NA | NA | NA | NA | Acp 62 | Acp 64 |
| | | U12 | L3 | HRM | NA | NA | NA | NA | Acp 62 | Acp 64 |
| | | U13 | L3 | HRM | NA | NA | NA | NA | Acp 62 | Acp 64 |
| | | U14 | L3 | HRM | NA | NA | NA | NA | Acp 62 | Acp 64 |
| | | U15 | L3 | HRM | NA | NA | NA | NA | Acp 62 | Acp 64 |
| | | U16 | L3 | HRM | NA | NA | NA | NA | Acp 62 | Acp 64 |
| | | U17 | L3 | HRM | NA | NA | NA | NA | Acp 62 | Acp 64 |
| | | U18 | L3 | HRM | NA | NA | NA | NA | Acp 62 | Acp 64 |
| | | U19 | L3 | PCR | NA | NA | NA | NA | Acp 62 | Acp 64 |
| | | U20 | L3 | PCR | NA | NA | NA | NA | Acp 62 | Acp 64 |
| | | U21 | L3 | PCR | NA | NA | NA | NA | Acp 62 | Acp 64 |

| Trio | Status | Label | MtDNA Lineage | Assignment method | Haplotype | Nuclear Cluster | Parentage assignment | | | |
|---|---|---|---|---|---|---|---|---|---|---|
| V | Adult | Acp70 | L3 | Seq | H16 | B | | | | |
| | | Acp72 | L3 | Seq | H17 | B | | | | |
| | | Acp66 | L1 | Seq | H7 | A | | | | |
| | | | | | | | Mother | Father | Parent 1 | Parent 2 |
| | Offspring | V1 | L3 | HRM+Seq | H16 | NA | Acp 70 | Acp 72 | NA | NA |
| | | V2 | L3 | HRM+Seq | H16 | NA | Acp 70 | Acp 72 | NA | NA |
| | | V3 | L3 | HRM+Seq | H16 | NA | Acp 70 | Acp 72 | NA | NA |
| | | V4 | L3 | HRM+Seq | H16 | NA | Acp 70 | Acp 72 | NA | NA |
| | | V5 | L3 | HRM+Seq | H16 | NA | Acp 70 | Acp 72 | NA | NA |
| | | V6 | L3 | HRM+Seq | H16 | NA | Acp 70 | Acp 72 | NA | NA |
| | | V7 | L3 | HRM+Seq | H16 | NA | Acp 70 | Acp 72 | NA | NA |
| | | V8 | L3 | HRM+Seq | H16 | NA | Acp 70 | Acp 72 | NA | NA |
| | | V9 | L3 | HRM+Seq | H17 | NA | Acp 72 | Acp 70 | NA | NA |
| | | V10 | L3 | HRM+Seq | H16 | NA | Acp 70 | Acp 72 | NA | NA |
| | | V11 | L3 | HRM+Seq | H17 | NA | Acp 72 | Acp 70 | NA | NA |
| | | V12 | L3 | HRM+Seq | H16 | NA | Acp 70 | Acp 72 | NA | NA |
| | | V13 | L3 | HRM+Seq | H16 | NA | Acp 70 | Acp 72 | NA | NA |
| | | V14 | L3 | HRM+Seq | H16 | NA | Acp 70 | Acp 72 | NA | NA |
| | | V15 | L3 | HRM+Seq | H17 | NA | Acp 72 | Acp 70 | NA | NA |
| | | V16 | L3 | HRM+Seq | H16 | NA | Acp 70 | Acp 72 | NA | NA |
| | | V17 | L3 | HRM+Seq | H16 | NA | Acp 70 | Acp 72 | NA | NA |
| | | V18 | L3 | HRM+Seq | H16 | NA | Acp 70 | Acp 72 | NA | NA |
| | | V19 | L3 | PCR+seq | H16 | NA | Acp 70 | Acp 72 | NA | NA |
| | | V20 | L3 | PCR+seq | H16 | NA | Acp 70 | Acp 72 | NA | NA |
| | | V21 | L3 | PCR+seq | H17 | NA | Acp 72 | Acp 70 | NA | NA |
| | | V22 | L3 | PCR+seq | H16 | NA | Acp 70 | Acp 72 | NA | NA |
| | | V23 | L3 | PCR+seq | H16 | NA | Acp 70 | Acp 72 | NA | NA |
